# Identification of P-type ATPase as a bacterial transporter for host-derived small RNA

**DOI:** 10.1101/2024.07.05.602285

**Authors:** Pu-Ting Dong, Mengdi Yang, Lujia Cen, Peng Zhou, Difei Xu, Peng Xiong, Chenggang Wu, Jiahe Li, Xuesong He

## Abstract

Host-derived small RNAs represent a finely tuned host response to maintain the host-microbial homeostasis. Among these, an emerging class known as transfer RNA-derived small RNA (tsRNA) has been implicated in modulating microbial-host interaction. Our previous study showed that when challenged with an oral pathobiont, *Fusobacterium nucleatum (Fn)*, an immortalized human oral keratinocyte cell line releases certain *Fn*-targeting tsRNAs that selectively inhibit the growth of *Fn* via their ribosome-targeting function. We also revealed the sequence- and energy-dependent uptake of tsRNAs by *Fn*. However, the mechanism underlying the tsRNA uptake at the molecular level remains elusive. In this study, using RNA affinity pull-down assay in combination with Mass Spectrometry, we identified a putative P-type ATPase transporter (PtaT) in *Fn*, which binds *Fn*-targeting tsRNAs in a sequence-specific manner. AlphaFold 3 simulation provides further evidence supporting the specific binding between PtaT and tsRNA compared to the scrambled control and the DNA counterpart. Through targeted mutagenesis and phenotypic characterization, we demonstrated the important role of PtaT in the uptake and antimicrobial capacity of tsRNAs against *Fn* in both ATCC 23726 type strain and a clinical tumor isolate (*Fn* CTI). Furthermore, global RNA sequencing and label-free Raman spectroscopy revealed the phenotypic differences between *Fn* wild type and PtaT-deficient mutant, highlighting the functional significance of PtaT in purine and pyrimidine metabolism. Collectively, our work identifies a bacterial transporter for tsRNAs and provides critical information for a fundamental understanding of how the host-generated tsRNAs specifically interact with its targeted bacteria at the molecular level.

## INTRODUCTION

Human mucosal surfaces provide a first line of protection against infectious bacteria through a complex array of innate and adaptive immunity^1,2^. The symbiotic relationship with hundreds of microbial species requires a finely tuned response at the mucosal surface to prevent the overgrowth of opportunistic pathogens while sparing the beneficial microbes^3–5^. Recent studies have highlighted certain processes, including host-derived small RNAs (sRNAs), that contribute to the maintenance of host-microbial homeostasis^6,7^. The intricate interplay between host-derived sRNAs and host-associated microbiome has emerged as a fascinating area of investigation, offering profound implications for understanding host-microbe interactions^8–11^.

Of particular interest are transfer RNA-derived small RNAs (tsRNAs) produced by endonucleases following the splicing of precursor or mature tRNAs^12^. tsRNAs have been shown to carry out important biological functions, such as epigenetic regulation, cell-cell communication, stress response and regulation of gene expression^12–16^, and can be aberrantly expressed in several diseased conditions^12,17^. Increasing lines of evidence also indicate that tsRNAs may play an important role in host-pathogen interactions^17^. Among human pathobionts, *Fusobacterium nucleatum* (*Fn*) represents a key player in various human diseases^18–21^, ranging from periodontitis^22,23^ to colorectal cancer^19,24,25^. Our previous study demonstrated that an immortalized human oral keratinocyte cell line releases two exosome-borne tsRNAs, tsRNA-000794 and tsRNA-020498 when challenged with *Fn* and these tsRNAs exhibit highly selective, *Fn*-targeting antimicrobial activity via their ribosome-targeting functions^26^. Chemical modification of these *Fn*-targeting tsRNAs, termed MOD-tsRNAs^26^, led to enhanced potency (over three orders of magnitude) while maintaining specificity^26^, thus offering promising prospects for targeted antimicrobial strategies. While our study revealed the sequence- and energy-dependent uptake of tsRNAs by *Fn*, the fundamental questions regarding the mechanisms underlying the cross-kingdom trafficking of host-generated *Fn*-targeting tsRNAs remain unanswered.

In the current study, we further elucidated the mechanistic underpinnings of the uptake of host-generated tsRNAs by *Fn* at the molecular level. By employing affinity pull-down, protein-RNA interaction simulation, coupled with targeted mutagenesis and bacteria phenotypic characterization, RNA-seq and Raman spectroscopy, we identified in *Fn* a putative tsRNA transporter, annotated as P-type ATPase transporter (PtaT). Our study demonstrated the important role of PtaT in mediating the uptake of host-generated tsRNAs and facilitating the antimicrobial activity of these *Fn*-targeting tsRNAs, thus providing a mechanistic and fundamental understanding of the tsRNAs-mediated cross-kingdom interaction.

## RESULTS

### Identification of a putative membrane protein binding to *Fn*-targeting tsRNAs

In previous studies^16,26^, we identified two host-derived *Fn*-targeting tsRNAs, tsRNA-000794 and tsRNA-020498, which are produced by an immortalized human oral keratinocyte cell line when challenged with *Fn*. These tsRNAs exhibit antimicrobial activity against *Fn* with high specificity. We further demonstrated sequence- and energy-dependent uptake of tsRNAs by *Fn*, however, it remains to be determined whether tsRNAs are internalized by *Fn* via an active transporting mechanism.

To query this, we sought to identify putative transporter proteins through the RNA affinity pulldown assay^27^, an established method for identifying sRNA-associated proteins in mammalian cells. Specifically, we prepared total bacterial lysate containing both cytoplasmic and membrane fractions, and used synthetic biotinylated tsRNA and streptavidin-conjugated magnetic beads to pull down putative proteins from the lysate that interact strongly with tsRNA-000794 and tsRNA-020498 but less so with the scrambled control, which could in principle enrich for target proteins (**Figure 1A**). As shown in the silver staining (**Figure 1B**), a band with a molecular weight of ∼75kDa was enriched in the samples from biotinylated tsRNA-000794 and tsRNA-020498 but noticeably less from the biotinylated scrambled RNA in three tested *Fn* strains (*Fn* ATCC 23726, *Fn* ATCC 25586 and *Fn* ATCC 10953) (**SI Fig. S1A**). Additionally, gel bands with the same molecular weight were detected in the tsRNA pulldown assays using six *Fn* clinical tumor isolates (CTIs) (**SI Fig. S1B**). In comparison, when the same RNA affinity pulldown assay was applied in total cell lysates from *Streptococcus mutans* (*Sm*) and *Porphyromonas gingivalis* (*Pg*), biotinylated tsRNA-000794 or tsRNA-020498 failed to enrich for any specific band compared to that of biotinylated scrambled RNA (**SI Fig. S1C**). Taken together, the presence of a unique protein band pulled down by two different *Fn*-targeting tsRNAs from *Fn* but not *Sm* or *Pg* may support the findings for species-specific tsRNA uptake as well as sequence-specific growth inhibition against *Fn*.

**Figure 1.**
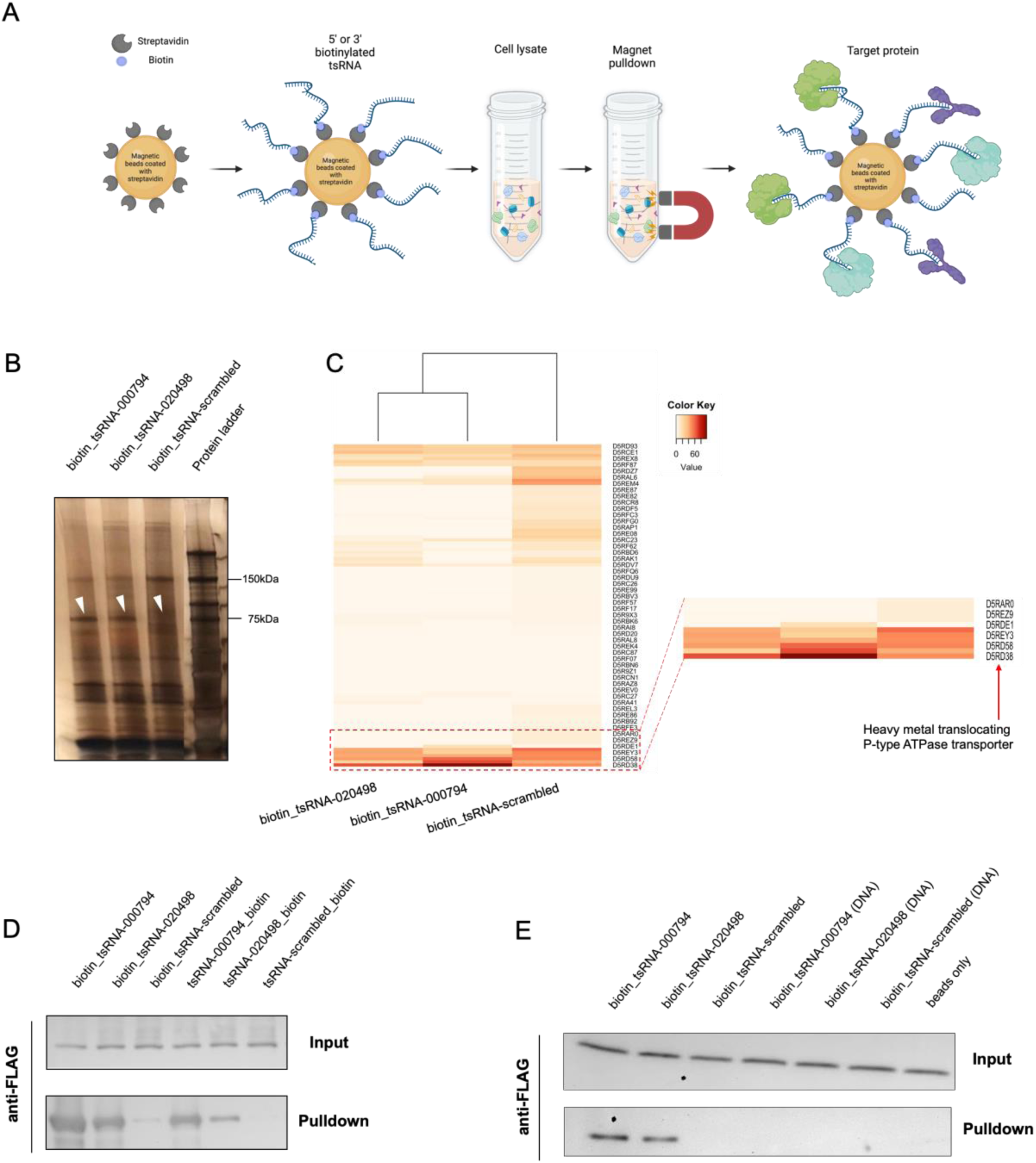
Identification of a putative tsRNA transporter protein in *Fn*. **A.** Illustration of the workflow for an RNA affinity pulldown assay to identify RNA-binding proteins. 5’ or 3’ biotinylated RNA oligonucleotides are immobilized on the surface of streptavidin-conjugated paramagnetic microparticles to capture RNA-interacting proteins from cell lysate. **B.** Silver staining of proteins from the biotinylated RNA pulldown in *Fn* ATCC 23726. Results are representative images of four independent experiments. **C.** Heatmap showing the relative abundance of proteins isolated from gel band at 75 kDa through Mass Spectrometry. The most significant band labeled as ‘D5RD38’ (P-type ATPase transporter, PtaT) was highlighted by a red arrow. **D.** Validating the interaction between biotinylated tsRNA and recombinant PtaT. 5’ or 3’ biotin tsRNA-mediated affinity pulldown was performed using the total lysate of *E. coli* BL21 Rosetta, in which FLAG-tagged PtaT was recombinantly expressed. 5’ and 3’ biotinylated scramble RNA serves as negative control. The upper panel (input) represents an equal amount of total lysates used, and the lower panel (pulldown) indicates the amount of FLAG-tagged PtaT specifically interacting with different biotinylated RNA. Representative image of two independent experiments is shown. **E.** Purified recombinant FLAG-PtaT can specifically bind tsRNA-000794 and tsRNA-020498 but not their DNA counterparts. Recombinant FLAG-tagged PtaT, which was purified from *E. coli* BL21 Rosetta, was directly used for the pulldown assay by 5’ biotinylated tsRNA. 5’ biotinylated scramble RNA and beads only served as negative control.

We then sought to identify the proteins specifically pulled down by the *Fn*-targeting tsRNAs via Mass Spectrometry. While certain RNases or known RNA-binding proteins were found in the gel bands at ∼75 kDa of molecular weight, they were not specifically enriched by the two *Fn*-targeting tsRNAs when compared to the scrambled control RNA (**Figure 1C**). In contrast, a putative membrane-bound P-type ATPase transporter (PtaT) was specifically associated with biotinylated tsRNA-000794 and tsRNA-020498 in *Fn* ATCC 23726, *Fn* ATCC 25586 and *Fn* ATCC 10953 as well as six *Fn* CTI strains. By aligning their amino acid sequences of PtaT, we found that ATCC 23726 and CTI-2 share identical protein sequences for PtaT, while other strains share 96-99% identities with ATCC 23726 (**Supplementary Table 1**). The conserved sequence for PtaT suggests that this protein may play a common role in tsRNAs binding, uptake, as well as tsRNA-mediated growth inhibition in tested *Fn* strains. However, since bioinformatic prediction suggested the function of PtaT in transporting metal ions, it remains unclear whether PtaT can bind and transport *Fn*-targeting tsRNAs.

To investigate the interaction between PtaT and tsRNAs, we ectopically expressed FLAG (DYKDDDDK) tagged PtaT (FLAG-PtaT) in *E. coli*. In agreement with the RNA affinity pull-down assay from the total lysate of *Fn* strains, 5’ or 3’-biotinylated tsRNA-000794 and tsRNA-020498 effectively pulled down FLAG-PtaT from the total lysate of *E. coli* overexpressing the target protein, while the scrambled control had minimal binding. To rule out the possibility that tsRNAs may indirectly bind FLAG-PtaT in the total lysates of *E. coli*, we further purified recombinant FLAG-PtaT from *E. coli* and demonstrated that biotinylated tsRNA-000794, and tsRNA-020498 but not the scrambled control RNA directly pulled down PtaT via streptavidin magnetic beads *in vitro* (**Figure 1D**). Additionally, we used purified FLAG-tagged PtaT along with magnetic beads conjugated with anti-FLAG antibodies as the bait to pull down tsRNA-000794, tsRNA-020498 or the scrambled control, and performed quantitative PCR to measure the levels of remaining free RNAs in the supernatant. As shown in **SI Fig. S2**, both tsRNAs exhibited interactions with the target proteins. Importantly, in this latter assay, the absence of biotin labeling, 2’ methylation and phosphorothioate bond in the two tsRNAs suggested that the direct binding between tsRNA and its target is dependent on the specific sequence rather than chemical modifications.

Having validated that PtaT can indeed interact with tsRNA-000794 and tsRNA-020498 of either chemically modified or naturally occurring ones, we next sought to examine whether PtaT was a bona-fide tsRNA-binding protein. To this end, we synthesized three 5’ biotinylated DNA oligos with identical chemical modifications, corresponding to tsRNA-000794, tsRNA-020498 and the scrambled RNA, respectively. We found that the recombinant *Fn* PtaT protein purified from *E. coli* can only bind tsRNA-000794 and tsRNA-020498 but not their DNA counterparts (**Figure 1E**). Meanwhile, we compared biotinylated tsRNA to DNA oligos of the same sequences by performing the same pulldown assays in *Fn* ATCC 23726 total lysate, followed by silver staining and Mass Spectrometry. Consistent with the direct binding experiment using purified PtaT, Mass Spectrometry results demonstrated tsRNA-000794 but not its DNA counterpart can pull down PtaT from *Fn* ATCC 23726 (**SI Fig. S3**). In addition to the scrambled control RNA, we found that two additional piwi-interacting RNAs (piRNAs) commonly found in human saliva did not pull down PtaT from *Fn* total lysate. In summary, through three different pulldown experiments including ectopic expression of FLAG-tagged PtaT in *E. coli* and *Fn*, respectively, as well as the use of purified PtaT, we showed that tsRNA-000794 and tsRNA-020498 can interact with PtaT in a highly sequence- and RNA-specific manner.

### AlphaFold 3 simulates the interaction between tsRNA-000794 and PtaT

To provide an additional line of evidence showing the interaction between *Fn*-targeting tsRNA and PtaT, we employed AlphaFold 3 to predict biomolecular interaction between PtaT from *Fn* ATCC 23726 and tsRNA-000794^28^. We downloaded the full protein sequence of PtaT for *Fn* ATCC 23726 from the UniProt website that corresponds to the protein identified by Mass Spectrometry for the tsRNA pulldown assay and chose tsRNA-000794 due to its higher uptake by *Fn* compared with tsRNA020498^26^. We then utilized AlphaFold 3 to predict the structure model of the tsRNA-000794 - PtaT protein complex (**Figure 2A**). The prediction results suggest that most regions of the protein exhibit high confidence, particularly the RNA-binding domain. Due to the limitations of current algorithms, the prediction confidence in tsRNA-000749 is lower than that of the protein. Nevertheless, the predicted protein-binding region in tsRNA still shows higher confidence compared to other regions (yellow), making it suitable for further analysis.

**Figure 2.**
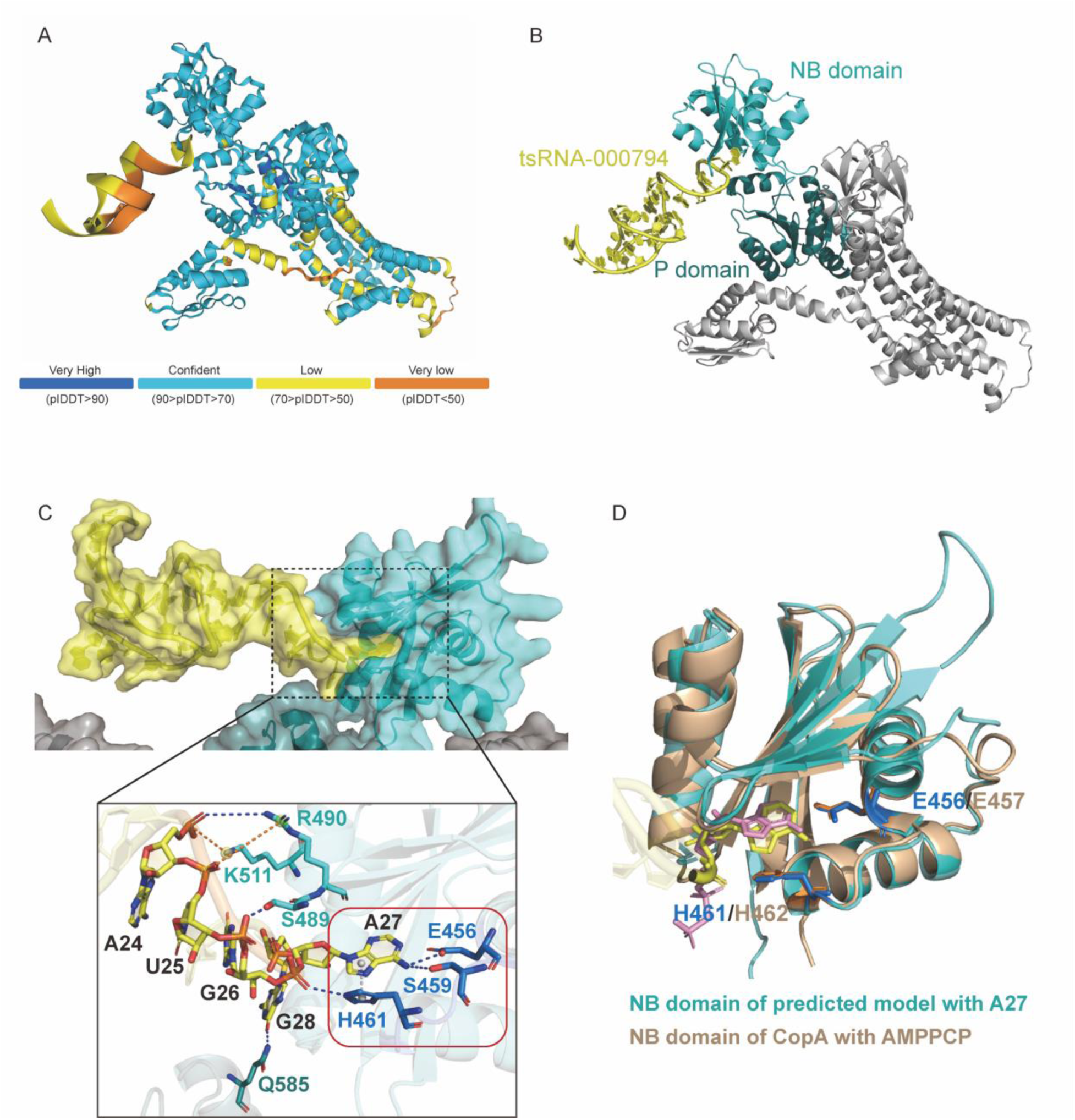
Structural analysis of the predicted complex model of tsRNA-000794 and PtaT. **A.** The predicted complex model of tsRNA-000794 and PtaT by AlphaFold 3. The color of the model presents the confidence of prediction, as shown in the bar below. **B.** Cartoon view of the predicted complex model. The tsRNA-000794 is shown in yellow, and the PtaT protein is shown in gray, with its Nucleotide Binding (NB) domain in cyan and Phosphorylation (P) domain in deep teal. **C.** Structural analysis of the binding pocket of the predicted model. Upper, binding region of tsRNA-000794 and PtaT. Both RNA and protein are shown in cartoon coated with surface view. Colors are the same as those in (**B**). Lower, detailed Interactions between tsRNA and NB domain/P domain of PtaT protein. Core pocket is highlighted in red brick. Nucleotide acids are in yellow. Key residues are shown in sticks. Residues in core pocket of NB domain are in blue, in extended region in NB domain are in cyan, and in P domain is in deep teal. Dotted lines represent interactions. Among them, gray indicates π-π interaction, blue indicates hydrogen bonding, and orange indicates salt bridge. **D**. Superimpose of the predicted model (cyan) and CopA (wheat, PDB code: 3A1C). Only NB domain is shown here for both proteins. A27 in tsRNA-000794 binding to core pocket of PtaT is in yellow stick, and AMPPCP, an ATP analog which binds to CopA is in pink stick. Key residues in core pocket also shown in stick, they are E456 H461 of PtaT in blue, and E457 H462 of CopA in orange. RMSD of NB domain between PtaT and CopA is 1.527Å.

Of note, P-type ATPase has a conserved Nucleotide Binding domain (NB domain), which is responsible for ATP binding and hydrolysis^29^. It also possesses a characteristic Phosphorylation domain (P domain), which is closely related to the NB domain and involved in the phosphorylation reaction following ATP hydrolysis^30^. Consistent with the above biochemical data, AlphaFold3 prediction suggested tsRNA-000794 may interact with *Fn* PtaT by contacting with both the NB domain and the P domain (**Figure 2B**).

A detailed analysis of the predicted PtaT-tsRNA interaction reveals a core pocket and an extended region in PtaT for tsRNA binding (**Figure 2C**). A27 of tsRNA is inserted into a core-binding pocket composed of residue E456-H461, T486, S489, G491 and V492 of PtaT. The interaction between A27 and E456, H461 and other pocket residues may stabilize the structure. For instance, in the depicted conformation (**Figure 2C**, lower), a T-shaped π-π interaction is formed between the sidechain of the purine ring of A27 and H461, while the -NH_2_ group on the purine ring of A27 forms hydrogen bonds with E456 and S459. E456 and H461 may play a role in the recognition and localization of A27, providing the specificity for tsRNA-000794 binding, and stabilizing A27 in the pocket. Thus, E456 and H461 are the key residues in the core pocket. In addition to the core pocket, a series of positively charged residues of PtaT, such as R490 and K511, are located along the direction of the tsRNA binding extension. These residues can form salt bridges with the phosphate backbone of the tsRNA. Furthermore, there are hydroxyl-containing amino acids, such as S489, which can form hydrogen bonds with the backbone of the tsRNA. Together, these residues form an extended region that helps maintain tsRNA binding. Moreover, G28 of tsRNA and residue Q585 in the P domain form hydrogen bonds in this model, which may further assist in binding.

The binding mode of the tsRNA-000794 to PtaT resembles the classical binding mode of ATP to P-type ATPases. For example, CopA proteins, a class of P-type ATPases that translocate Cu^2+^, have had their crystal structures solved, including those containing the ATP analogue - AMPPCP^31^. The CopA - AMPPCP complex structure (PDB code: 3A1C) can be used as a model to study ATP binding to P-type ATPases, serving as the homologue to the PtaT protein. The superimposition of the NB domains of the PtaT protein and CopA reveals a high degree of similarity, especially at the core pocket site and key residues (E456 and H461 in PtaT protein, E457 and H462 in CopA, respectively). Correspondingly, A27 of tsRNA-000794 exhibits a similar pose inserted into the core pocket as AMPPCP (**Figure 2D**).

We further predicted the interaction between *Fn* PtaT and tsRNA scrambled control, or the DNA counterpart of tsRNA-000794 (tsDNA-000794). As shown in **SI Fig. S4A**, the scrambled control appears to adopt a relatively stable conformation within the complex. However, it lacks the specificity to bind to the pocket in the NB domain, as seen with the tsRNA-000794. The scrambled control may form electrostatic interactions with K562 and R566, potentially leading to binding in the P domain. Nevertheless, this binding does not exhibit specific binding property as that observed with tsRNA-000794. We also simulated the interaction between tsDNA-000794 and PtaT, and we found no binding between them (**SI Fig. S4B**).

In summary, the predicted structure model of tsRNA-000794 - PtaT protein complex by AlphaFold 3 is referable, as evidenced by both structural analysis and homologous comparison. Additionally, the predicted binding mode provides insights into the binding affinity and specificity of tsRNA-000794 to PtaT, which agrees with the biochemical data.

### Knocking out *ptaT* interferes with the intake and antimicrobial efficacy of *Fn*-targeting tsRNAs

Having demonstrated that PtaT is a possible RNA binding protein for tsRNA-000794 and tsRNA-020498 experimentally and computationally, we next explored the functions of PtaT in tsRNA-mediated growth inhibition of *Fn*. Here, we anticipated two possible roles for PtaT: 1) If it is involved in tsRNA uptake, or processing, knocking out *ptaT* will confer resistance against tsRNA-mediated *Fn* inhibition; 2) if it is the target of tsRNAs whose function is critical for *Fn* physiology and can be directly inhibited by tsRNAs, *Fn* mutant defective in *ptaT* may display a growth defect or may not even be viable.

To knock out *ptaT* in *Fn* 23726, generating a *galK* mutant strain in *Fn* ATCC 23726 (Δ*galK*) is often a first step by harnessing the *galK* gene as a counter-selectable marker^32,33^. Specifically, the absence of *galK* allows for the selection of mutants when grown on specific media that exploit the galactose metabolism pathway^34^. For example, mutants lacking *galK* can survive on media containing 2-deoxy-galactose (2-DG), which is toxic to cells that possess an active *galK* gene. After obtaining the *Fn* ATCC 23726 Δ*galK* mutant, a suicide vector was designed to enable a double crossover mediated removal of the *ptaT* gene in the Δ*galK* strain background, and the successful deletion of both *galK* and *ptaT* genes was verified by colony PCR and Sanger sequencing (**SI Fig. S5**), resulting in *Fn* Δ*galK* Δ*ptaT* mutant.

We first observed that *Fn* Δ*galK* and *Fn* Δ*galK* Δ*ptaT* grew similarly in the Columbia Broth media (**SI Fig. S6**), suggesting that deletion of *ptaT* doesn’t significantly affect the bacterial viability under the standard culturing condition. Next, we treated *Fn* Δ*galK* and *Fn* Δ*galK* Δ*ptaT* with chemically modified tsRNA-000794 (MOD-(OMe)-000794) and the scrambled control (MOD-(OMe)-scrambled), respectively.

We then examined the antimicrobial efficacy of tsRNAs on *Fn* through a SYTOX Green assay described previously^26,35^. As expected, MOD-(OMe)-000794 induced significant cell death in *Fn* Δ*galK*, indicated by a large portion of cells exhibiting green fluorescence (**Figure 3A**). In comparison, *Fn* Δ*galK* Δ*ptaT* was relatively resistant to tsRNA-mediated growth inhibition compared to *Fn* Δ*galK* (**Figure 3A**). MOD-(OMe)-scrambled control didn’t exert an inhibitory effect towards both *Fn* Δ*galK* and *Fn* Δ*galK* Δ*ptaT* (**Figure 3B**), as evidenced by very few SYTOX Green-positive bacteria. Quantification of integrated SYTOX Green fluorescence signal after normalization of the total bacterial area further underscored that PtaT indeed plays an important role in mediating tsRNA-induced growth inhibition of *Fn* (**Figure 3C**).

**Figure 3.**
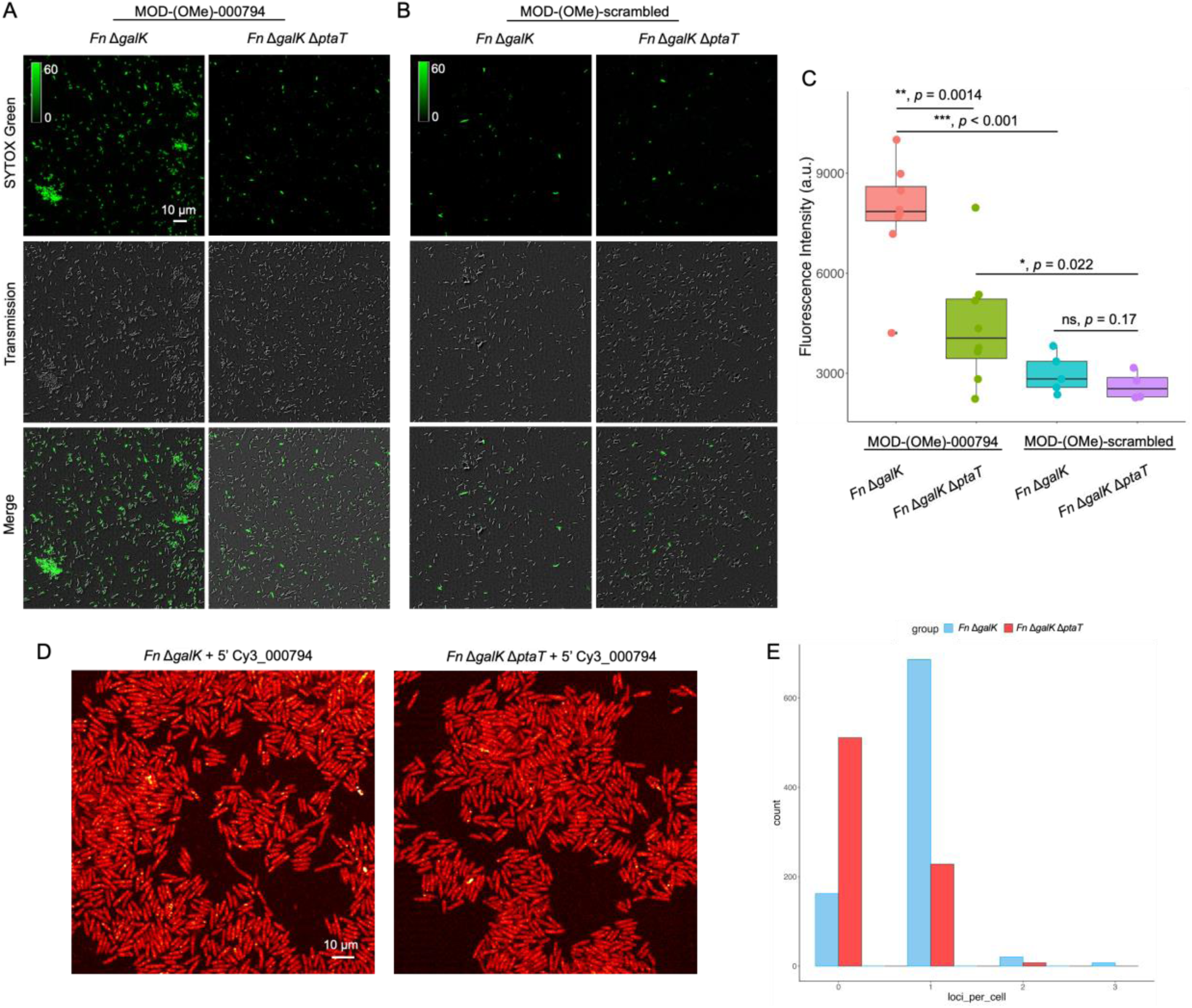
P-type ATPase transporter (PtaT) plays an important role in the internalization and antimicrobial effect of modified tsRNAs on *Fusobacterium nucleatum* ATCC 23726. **A-B**. *Fn* Δ*galK* and *Fn* Δ*galK* Δ*ptaT* were treated with 500 nM MOD-(OMe)-000794 (**A**), and MOD-(OMe)-scrambled (**B**) for 5 hours followed by SYTOX Green staining. **C**. SYTOX Green quantification was carried out by normalizing raw integrated fluorescence intensity to the areas of randomly picked bacteria, which takes into consideration of both SYTOX Green positive and -negative ones in the field of view. Data were representative of three independent experiments and analyzed by the Student’s unpaired *t*-test. ** *p* < 0.01, *** *p* < 0.001. **D**. Visualization of internalization of 5’ Cy3_000794 by *Fn* Δ*galK* and *Fn* Δ*galK* Δ*ptaT* through super-resolution Airyscan confocal microscopy. *Fn* Δ*galK* and *Fn* Δ*galK* Δ*ptaT* were incubated with 5’ Cy3_000794 for 5 hours followed by imaging. **E**. Histogram showing the number of loci (localized dense accumulation of 5’ Cy3_000794) per cell between *Fn* Δ*galK* and *Fn* Δ*galK* Δ*ptaT*.

To query whether the increased resistance to tsRNA in *Fn* Δ*galK* Δ*ptaT* is due to the reduced tsRNA intake, we fluorescently labeled the tsRNAs with Cy3 at the 5’ end, and then treated both *Fn* Δ*galK* and *Fn* Δ*galK* Δ*ptaT* with Cy3-tagged tsRNA-000794. Super-resolution Airyscanning fluorescence microscopy was then applied to examine the intracellular accumulation of tsRNA. As shown in **Figure 3D**, Cy3-tagged tsRNA-000794 indeed was taken up by *Fn* Δ*galK*, resulting in intracellular fluorescence signal with the formation of ‘bright dots’ (loci) indicative of subcellular accumulation of tsRNA. However, by knocking out *ptaT*, the intracellular accumulation of Cy3-tagged tsRNA-000794 was largely reduced. This difference was documented by the distribution disparity of the number of loci per bacteria between these two groups (**Figure 3E**). Taken together, these data provide genetic and phenotypic evidence on the roles of PtaT in tsRNA uptake and growth inhibition in *Fn*.

### Knocking out *ptaT* affects the global RNA profiles of *Fn*

Our data so far demonstrated the role of PtaT in mediating the inhibitory effect of host-derived tsRNAs against *Fn*. To further investigate the additional biological role of PtaT in *Fn*, we performed RNA-seq to identify the differentially expressed genes (DEGs) in *Fn* Δ*galK* Δ*ptaT* compared to *Fn* Δ*galK* in both log and early stationary phases.

Gene expression levels were compared between *Fn* Δ*galK* and *Fn* Δ*galK* Δ*ptaT* during log-phase growth in three biological replicates. A total of 580 DEGs with a false discovery rate (FDR)-adjusted *p*-value < 0.05 are identified and presented by heatmaps and volcano plot (**Figure 4**). KEGG enrichment scatter plot of DEGs (**Figure 4A**) and quantification of differentially expressed genes analysis (**Figure 4B**) showed that purine metabolism represents one of the most significantly downregulated pathways in *Fn* Δ*galK* Δ*ptaT* (**Figure 4, A-B**). Other significantly downregulated pathways include biosynthesis of secondary metabolites; pyrimidine metabolism, pyruvate metabolism; alanine, aspartate and glutamate metabolism (**Figure 4E**). Log-phase *Fn* Δ*galK* Δ*ptaT* also displayed increased expression of glycerophosphoryl diester phosphodiesterase and genes related to methionine metabolism (**Figure 4E**).

**Figure 4.**
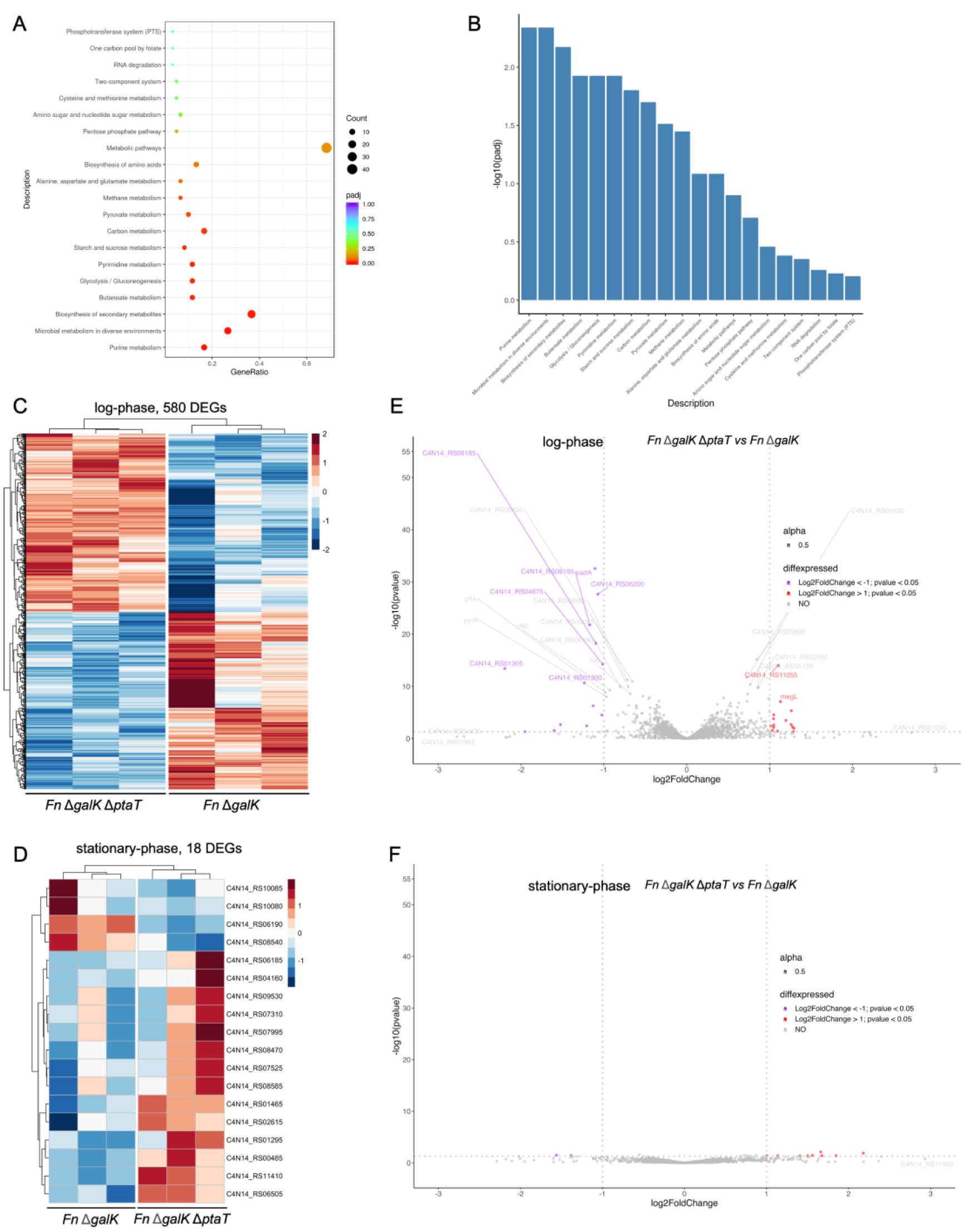
Transcriptomic analysis of log-phase and stationary-phase *Fn* Δ*galK* and *Fn* Δ*galK* Δ*ptaT*. **A-B**. Clusters of orthologous groups (COG, **A**) and quantification of differentially expressed genes (**B**) from log-phase *Fn* Δ*galK* Δ*ptaT* relative to *Fn* Δ*galK*. **C-D**. Heatmap showing the global differentially expressed genes for both log-phase (**C**) and stationary-phase (**D**) *Fn* Δ*galK* and *Fn* Δ*galK* Δ*ptaT.* Each heatmap includes triplicate RNA-seq samples for the indicated *Fn* Δ*galK* and *Fn* Δ*galK* Δ*ptaT*. The coloring indicates Log2FoldChange of the selected samples, while red and blue denote up- and down-regulation, respectively. The DESeq2 method (*p*-value <= 0.05) was applied to generate the heatmap. **E**-**F**. Volcano plots showing transcriptomic changes of log-phase (**E**) and stationary-phase (**F**) *Fn ΔgalK ΔptaT* relative to *Fn* Δ*galK.* Red and magenta dots indicate significantly up-regulated and down-regulated genes, respectively, and grey dots indicate genes with no significant changes. Significantly differentially regulated genes are characterized by an absolute fold change >2 (down-regulated log2 < −1, up-regulated log2 > 1; vertical dashed line) and *p*-value < 0.05 (horizontal dashed line).

On the contrary, only 18 DEGs were identified between *Fn* Δ*galK* and *Fn* Δ*galK* Δ*ptaT* during stationary-phase growth (**Figure 4D**). As shown in the volcano plot (**Figure 4F**) and the clusters of orthologous group plot (**SI Fig. 7**), two DEGs displayed reduced expression in *Fn* Δ*galK* Δ*ptaT* compared to *Fn* Δ*galK*: C4N14_RS10085 (related to glutamine metabolism); C4N14_RS10080 (related to carbamoyl-phosphate metabolism). Nine DEGs showed upregulated gene expression which are related to lipopolysaccharide biosynthesis (C4N14_RS09530); tRNA activity (C4N14_RS01465), and histidine phosphatase (C4N14_RS08585). The drastic difference in gene expression pattern between *Fn* Δ*galK* Δ*ptaT* and *Fn* Δ*galK* in log-phase and stationary-phase bacteria echoed with what has been reported previously: the metabolism-linked genes are highly expressed when cells are growing and get turned off when the cells enter stationary phase^36–38^. Taken together, the transcriptomic data highlight the significant role that PtaT plays in shaping the global metabolic profiles of *Fn,* particularly during its actively growing state.

### Raman spectroscopy revealed the PtaT-dependent global metabolic profiles of *Fn*

Purine and pyrimidine are involved with the major energy carriers, and they are the subunits of nucleic acids^39,40^. Since our transcriptomic data indicates potentially impaired purine synthesis in *Fn* Δ*galK* Δ*ptaT* in the log phase, we wondered whether the reduced gene expression of purine synthesis-related genes in the log phase would later result in a lower abundance of nucleic acids inside bacteria. To answer this question, we first analyzed and compared bulk RNA levels in the stationary-phase cells. It was found that the level of bulk RNA extracted from an equal amount of *Fn* Δ*galK* was ∼two-fold higher than that of *Fn* Δ*galK* Δ*ptaT* (*p* = 0.037, **SI Fig. S8**). Interestingly, there was no significant difference between log-phase *Fn* Δ*galK* Δ*ptaT* and *Fn* Δ*galK* in the total RNA levels (*p* = 0.77). These findings suggested a delayed response at the RNA levels following decreased gene expression related to purine/nucleic acid metabolism. To further corroborate the difference in total bulk RNA extraction experiments, we performed Raman spectroscopy to understand how PtaT-depletion impacted *Fn* Δ*galK* at the molecular and cellular levels. Raman spectroscopy has been widely used to provide insights into the chemical makeup of biological samples at single-cell level^41–43^, and we sought to obtain spectroscopic vibrational information of intracellular biomolecules from both the log- and stationary-phase *Fn* Δ*galK* and *Fn* Δ*galK* Δ*ptaT.* As shown in **Figure 5A-B**, both log- and stationary-phase *Fn* Δ*galK* and *Fn* Δ*galK* Δ*ptaT* exhibited typical Raman peaks: 720/780 cm^-1^ (DNA/RNA); 1003 cm^-1^ (phenylalanine); 1240 cm^-1^/1450 cm^-1^/1660 cm^-1^ (Amide III/II/I peaks).

**Figure 5.**
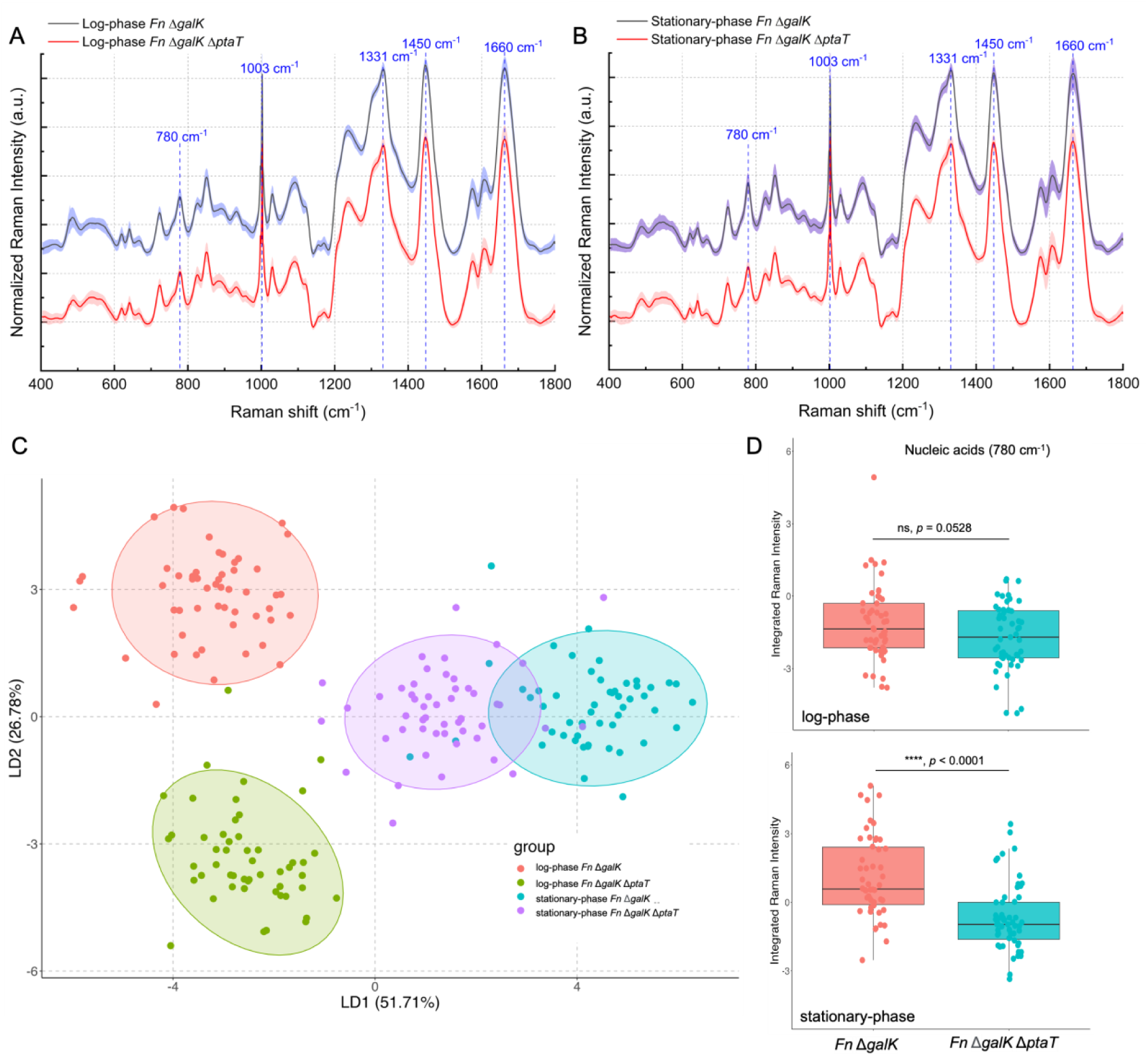
Spectroscopic characterization of both log-phase and stationary-phase *Fn* Δ*galK* and *Fn* Δ*galK* Δ*ptaT* by label-free Raman spectroscopy. **A-B.** Averaged Raman spectra of log-phase (**A**) and stationary-phase (**B**) *Fn* Δ*galK* and *Fn* Δ*galK* Δ*ptaT.* Each scenario was averaged from 50 spectra over three biological replicates. Shaded area represents standard deviation from the measurements. Peaks of interest were highlighted by dashed blue lines and designated with corresponding Raman shifts. **C**. Linear discrimination analysis (LDA) of 200 Raman spectra from log-phase and stationary-phase *Fn* Δ*galK* and *Fn* Δ*galK* Δ*ptaT.* Plot was shown through the display of LD2 versus LD1. 95% confidence intervals were outlined by colored ellipses. **D.** Quantification of the amount of nucleic acids from both log-phase and stationary-phase *Fn* Δ*galK* and *Fn* Δ*galK* Δ*ptaT.* Quantification was calculated through the integration of Raman intensity from 770 cm^-1^ to 789 cm^-1^. Statistical analysis was achieved by the Student’s unpaired *t*-test: **** *p* < 0.0001, ns, not significant.

To globally map out the difference among the four groups, a multivariate data analysis approach, linear discrimination analysis^44,45^ (LDA) was utilized to model the difference among the four groups through dimensionality reduction of high-dimension Raman spectra. Each group indeed exhibited a distinct cluster in the LDA plots (**Figure 5C** and **SI Fig. S9**). Raman band at 780 cm^-1^ is related to nucleic acids due to the cytosine/uracil ring breathing^46^. We then quantified the amount of nucleic acids based on the integrated Raman intensity around 780 cm^-1^ given that Raman intensity is linearly proportional to the amount of biomolecules inside the samples^47^. We found that *Fn* Δ*galK* Δ*ptaT* had significantly lower nucleic acid levels (*p* < 0.0001) than that of the isogenic control *Fn ΔgalK* in the stationary phase compared to that of the log phase (ns, *p* = 0.0528), suggesting that the reduced purine synthesis likely limited nucleic acids level in the later stage (**Figure 5D**). Consistent with the transcriptomic data and RNA quantification from the bulk sample (**SI Fig. S8**), label-free Raman spectroscopy provides another line of evidence that PtaT may affect the purine synthesis process which negatively impacts intracellular nucleic acids level at the stationary phase.

### Knocking out *ptaT* in a *Fn* clinical tumor isolate interferes with the intake and antimicrobial efficacy of *Fn*-targeting tsRNAs

After characterizing the roles of PtaT in tsRNA’s uptake, tsRNA-mediated growth inhibition and purine metabolism in the type strain *Fn* ATCC 23726, we next explored if similar findings can be validated in a clinical *Fn* isolate. *Fn* is a significant contributor to colorectal cancer (CRC)^48^. Since we have shown the excellent antimicrobial efficacy of MOD-(OMe)-000794 against multiple *Fn* clinical tumor isolates (CTIs)^26^, we wondered whether PtaT plays similar roles in tsRNA intake and its induced growth inhibition as observed in *Fn* ATCC 23726.

To test this, we chose *Fn* clinical tumor isolate (CTI)-2 because CTI-2 expresses a membrane protein, Fap2 that mediates the adhesion of *Fn* to colon cancer cells overexpressing Gal-GalNAc to promote the development of CRC^49^. Additionally, we also found that the protein sequences for PtaT are identical between CTI-2 and *Fn* ATCC 23726. For these reasons, we generated a PtaT-depleted *Fn* CTI-2 strain (*Fn* CTI Δ*ptaT*) as described in Materials and Methods. After validating the successful knockout of *ptaT*, we examined the inhibition efficacy of tsRNAs on *Fn* CTI-2 wildtype (*Fn* CTI wt) and *Fn* CTI Δ*ptaT* through the SYTOX Green assay. MOD-(OMe)-000794 induced apparent cell death in the case of *Fn* CTI wt, indicated by a large portion of cells exhibiting green fluorescence (**Figure 6A**) and consistent with our previous study^26^. Similar to findings in *Fn* 23726 Δ*galK* Δ*ptaT*, we also observed that *Fn* CTI Δ*ptaT* was relatively resistant to tsRNA-mediated growth inhibition compared to *Fn* CTI wt (**Figure 6A**). MOD-(OMe)-scrambled didn’t exert any inhibitory effect towards both *Fn* CTI wt and *Fn* CTI Δ*ptaT* (**Figure 6B**), as evidenced by few SYTOX Green-positive bacteria. Quantification of integrated SYTOX Green fluorescence signal after normalization of the total bacterial area further underscored that PtaT indeed plays an indispensable role in mediating tsRNA-mediated growth inhibition of a *Fn* clinical tumor isolate (**Figure 6C**).

**Figure 6.**
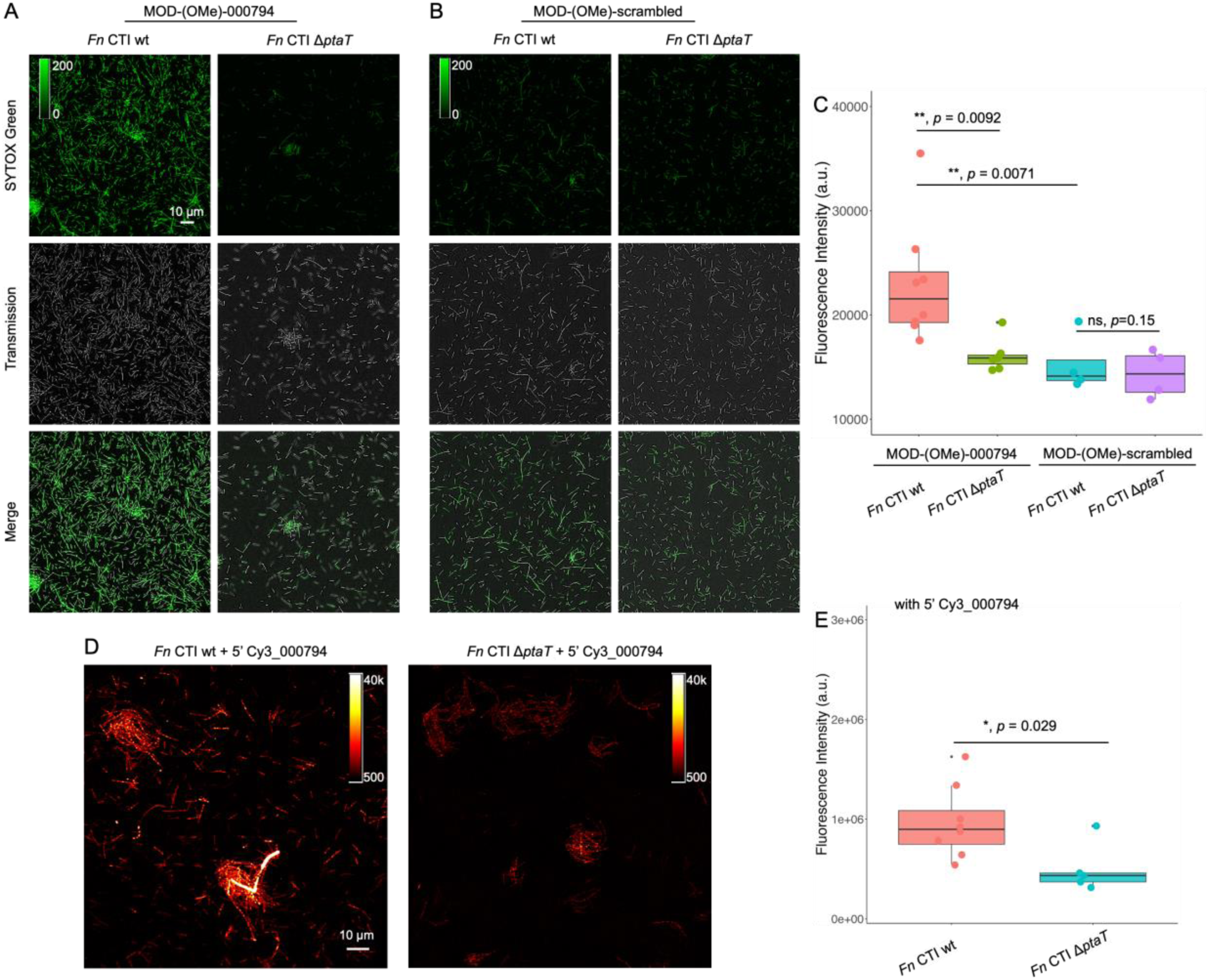
P-type ATPase transporter (PtaT) plays an essential role in the internalization and antimicrobial effect of modified tsRNAs on a *Fusobacterium nucleatum* clinical tumor isolate (*Fn* CTI). **A-B**. *Fn* CTI wt and *Fn* CTI Δ*ptaT* were treated with 500 nM MOD-(OMe)-000794 (**A**), and MOD-(OMe)-scrambled (**B**) for 5 hours followed by SYTOX Green staining. **C**. SYTOX Green quantification was carried out by normalizing raw integrated fluorescence intensity to the areas of randomly picked bacteria, which takes into consideration of both SYTOX Green positive and negative ones in the field of view. Statistical analysis was achieved by the Student’s unpaired *t*-test: ** *p* < 0.01, * *p* < 0.05, ns, not significant. **D**. Visualization of internalization of 5’ Cy3_000794 by *Fn* CTI wt and *Fn* CTI Δ*ptaT* through confocal fluorescence imaging. *Fn* CTI wt and *Fn* CTI Δ*ptaT* were incubated with 5’ Cy3_000794 for 5 hours followed by confocal fluorescence imaging. **E**. Quantification of fluorescence intensity by normalizing raw integrated fluorescence intensity to the areas of bacteria as shown **D**. Statistical analysis was achieved by the Student’s unpaired *t*-test: * *p* < 0.05.

To investigate if the increased resistance to tsRNA in *Fn* CTI Δ*ptaT* is due to the reduced tsRNA intake, we treated both *Fn* CTI wt and *Fn* CTI Δ*ptaT* with 5’-Cy3-tagged tsRNA-000794. Confocal fluorescence microscopy was then applied to examine the intracellular accumulation of tsRNA. As shown in **Figure 6D**, Cy3-tagged tsRNA-000794 was densely accumulated intracellularly as indicated by the formation of multiple loci with strong signal intensity. By knocking out *ptaT*, the intracellular accumulation of Cy3-tagged tsRNA-000794 was significantly reduced in the *Fn* CTI Δ*ptaT* compared to *Fn CTI* wt (*p* = 0.029, **Figure 6E**). This difference was documented by the comparison of the integrated Cy3 fluorescence intensity after normalization by dividing the total bacterial area. These data provide further evidence showing the important role of PtaT in the uptake of tsRNA and its mediated growth inhibition against a wide range of *Fn* strains, including clinically relevant ones.

## DISCUSSION

Small regulatory noncoding RNAs (sRNAs), including transfer RNA-derived sRNAs (tsRNAs), are a class of regulatory elements that have been identified in prokaryotes^50^ and eukaryotes^51^ and implicated in gene regulation. Recent studies further revealed the potential role of sRNAs in interspecies and cross-domain interactions^52^ with increasing lines of evidence showing that host-derived sRNAs such as fecal miRNAs can modulate the compositions of host-associated microbiota^53,54^.

Over the last decades, transmembrane RNA importer proteins in eukaryotes, such as Systemic RNA Interference Deficiency-1 (SID-1)^55^ in *Caenorhabditis elegans*, and its homologs SIDT1^56^ and SIDT2^57^ in mammals, have been identified that facilitate internalization of extracellular sRNAs for intercellular or cross-kingdom gene modulation. However, little progress has been made toward elucidating how host-derived sRNAs may enter bacteria in a cross-kingdom fashion. In this work, through comprehensive biochemical analysis, we identified a P-type ATPase transporter (named PtaT), a putative membrane-associated protein in *Fn* that is capable of binding tsRNAs in a sequence-dependent manner. More importantly, our genetic and phenotypic data strongly support PtaT’s role in mediating the uptake and growth inhibition of *Fn*-targeting tsRNAs. It is worth noting that while deletion of *ptaT* in *Fn* led to a significant decrease in the uptake of and reduced sensitivity to *Fn*-targeting tsRNAs, it did not completely abolish the intake of tsRNAs, suggesting the presence of other yet-to-be-identified cellular function(s) that may also contribute to tsRNAs uptake.

P-type ATPases are a large group of evolutionarily related integral membrane proteins found in bacteria, archaea and eukaryotes. Most of the characterized P-type ATPases are active pumps that couple ATP hydrolysis to the transport of diverse substrates, ranging from H^+^, metal cations, phospholipids to polyamines^58,59^. A better understanding of the potential role of PtaT in transporting *Fn*-targeting tsRNA will benefit from the detailed structural characterization of PtaT-tsRNA complex using X-ray crystallography or cryo-electron microscopy. Nevertheless, our data strongly indicate that PtaP in *Fn* may bind to exogenous sRNAs and facilitate their intake, thus likely expanding the substrate diversity of P-type ATPase transporter in bacteria.

While *ptaT*-depleted *Fn* mutant does not display significant growth defects compared to *Fn* wt under optimal growth conditions, a significantly downregulated expression in purine metabolism-related genes was observed during its log-phase growth, followed by a drastic reduction in intracellular nucleic acids level at the stationary phase. Interestingly, our previous study^26^ also revealed that treatment of *Fn* with *Fn*-targeting tsRNA markedly downregulated the same purine metabolism pathway compared to the scrambled control. While further investigation is warranted, it is tempting to speculate that PtaT may contribute to purine biosynthesis by facilitating the uptake of precursor molecules involved with purine biosynthesis. This function could be negatively impacted as a result of the competitive binding of PtaT by *Fn*-targeting tsRNAs. While it is to be determined if *Fn* PtaT could bind to multiple substrates, studies of P-type ATPase ATP12A3 suggested that it may bind to both K^+60^ and polyamine^61^ to mediate their co-transportation or K^+^ and polyamines may compete for the same binding sites.

Interestingly, PtaT protein shares high protein sequence identities (96-100%) among three *Fn* type strains and six clinical isolates that we characterized in this study. It is also highly conserved (∼95% amino acid identities) within the *Fusobacterium* genus. In contrast, the most homologous proteins found in *Sm* ATCC 6249, *Pg* ATCC 33277 and *E. coli* MG1655 are only 31%, 30% and 27% identical to PtaT from *Fn* ATCC 23726. This agrees with our previous finding that, while *Fn*-targeting tsRNAs can be found to bind to the cell surface of *Pg*, they failed to be internalized^26^, suggesting that sequence-dependent binding between PtaT and tsRNA may contribute to the observed tsRNA-mediated antimicrobial specificity. Thus, understanding the role of host-derived tsRNA uptake mechanisms in *Fn* pathogenesis may have far-reaching implications for developing targeted therapeutics and interventions against *Fn*-associated diseases.

Given the abundance of extracellular host-derived sRNAs and their suspected role in microbial-host interaction, it would be intriguing to explore whether PtaT is a sRNA-transporter unique to *Fn*, or P-type ATPases may represent a more generalized portal through which host-derived tsRNAs may be selectively imported into bacteria. By identifying a novel bacterial transporter for host-derived small RNA, our study offers novel insights into sRNA-mediated host-pathogen interplay in the context of microbiome-host interactions.

## Materials and Methods

### Chemicals

All chemicals and cell culture broth were purchased from Fisher Scientific International Inc. (Cambridge, MA, USA) unless otherwise noted, and were of the highest purity or analytical grade commercially available. DNA and RNA oligos were ordered from Sigma Aldrich (St. Louis, MO, USA) and Integrated DNA Technologies (Coralville, IA, USA). All molecular cloning reagents including restriction enzymes, competent cells, and the Gibson assembly kit were purchased from New England Biolabs (Ipswich, MA, USA) .

### Bacterial strains (summarized in **Table 1**) and growth conditions

*Fusobacterium nucleatum subsp. nucleatum* ATCC 23726, 25586, 10953, *Streptococcus mitis* ATCC 6249, and *Porphyromonas gingivalis* ATCC 33277 were purchased from the American Type Culture Collection (Manassas, VA, USA). *F. nucleatum* colon tumor isolate was general gift of Dr. Wendy Garrett at the Harvard T.H. Chan School of Public Health. *F. nucleatum* strains and *P. gingivalis* were cultured in liquid Columbia broth (CB) or on CB agar plates containing 5% defibrinated sheep blood, and incubated at 37°C in an anaerobic chamber (Sheldon Manufacturing, Cornelius, OR, USA) containing 5% H_2_, 10% CO_2_, 85% N_2_. *S. mitis* was cultured in Brain-Heart Infusion (BHI) broth. *E. coli* was cultured in lysogeny broth (LB) media and incubated at 37°C under aerobic condition. Thiamphenicol at 5 μg/mL (Fisher Scientific) was used for the selection and maintenance of *Fn* strains possessing pHS31. *F. nucleatum* CTI-2 strain and its derivatives were cultured in a TSPC medium comprising 3% tryptic soy broth (BD), 1% Bacto peptone, and 0.05% cysteine. Fusobacterial transformants with pBCG02-based deletion plasmid were grown overnight in tryptic soy broth supplemented with 1% Bacto peptone plus 0.25% freshly made cysteine (TSPC) broth with 5 µg/mL thiamphenicol. *Escherichia coli* strains were grown in Luria-Bertani (LB) broth with aeration at 37 °C. *E. coli* strains carrying plasmids were grown in LB broth containing 20 µg/mL chloramphenicol.

**Table 1.** Bacterial strains and plasmids used in this paper.

| Bacterial Species | Strains | Characteristics | Source |
| --- | --- | --- | --- |
| *Porphyromonas gingivalis* | ATCC 33277 | WT | ATCC |
| *Streptococcus mitis* | ATCC 6249 | WT | ATCC |
| *F. nucleatum* | ATCC 23726 | *ssp. nucleatum* WT | ATCC |
|  | ATCC 25586 | *ssp. nucleatum* WT | ATCC |
|  | ATCC 10953 | *ssp. nucleatum* WT | ATCC |
|  | *Fn_ΔgalK* | *galK* markerless deletion | this study |
|  | *Fn_ΔgalK ΔptaT* | *ptaT* markerless deletion | this study |
|  | CTI-1 (*ssp. animalis*) | clinical tumor isolate | Previous study<sup>49</sup> |
|  | CTI-2 (*ssp. nucleatum*) | clinical tumor isolate | Previous study<sup>49</sup> |
|  | CTI-3 (*ssp. animalis*) | clinical tumor isolate | Previous study<sup>49</sup> |
|  | CTI-5 (*ssp. animalis*) | clinical tumor isolate | Previous study<sup>49</sup> |
|  | CTI-6 (*ssp. polymorphum*) | clinical tumor isolate | Previous study<sup>49</sup> |
|  | CTI-7 (*ssp. vincentii*) | clinical tumor isolate | Previous study<sup>49</sup> |
|  | CTI-2 *ΔptaT* | deletion of *ptaT* in CTI-2 | this study |
| *Escherichia coli* | NEB5α |  | NEB |
|  | Rosetta (DE3) |  | NEB |
|  | C2987 | cloning host | NEB |
| **Plasmids** | **Purpose** | **Characteristics** | **Source** |
| pSH200-PtaT | PtaT purification | amp | this study |
| pHS31_FLAG_ *galK* | In-frame deletion of *galK* | *catP* | this study |
| pHS31_FLAG-*galK*-Δ*ptaT* | In-frame deletion of *ptaT* | *catP* | this study |
| pBCG02 | *E. coli*/*Fusobacterium* shuttle vector | *cm<sup>R</sup>/thia<sup>R</sup>* | ref<sup>32</sup> |
| pBCG02-CTI2-1787updn | Derivative of pBCG02 lacking *repA* and *ori<sub>Fn</sub>*, deletion plasmid of *ptaT* |  | this study |

### Plasmid construction for genetic knockout of F. nucleatum ATCC 23726

A list of primers is provided in **Table 2**. For expression and purification of recombinant PtaT in *E. coli*, the full-length cDNA was amplified from the genomic DNA of *F. nucleatum* ATCC 23726 genome via primers B267/B268 and ligated to pSH200 via *BamH I* and *Not I* by the Gibson Assembly kit. His-tagged proteins were expressed in Rosetta (DE3) *E. coli*. For constructing a suicide vector for insertional mutagenesis of PtaT in *Fn*, a 1kb central fragment was cloned into pHS31 after *SnaBI* digestion through the Gibson assembly. For overexpression in *Fn*, a shuttle vector, pHS58 was first digested with *XhoI* and *HindIII*. The FN1529 promoter amplified from *Fn 25586*, and the full-length *ptaT* were assembled with digested pHS58. All constructs were first cloned into NEB 5-alpha chemically competent *E. coli*, and then verified by Sanger sequencing before transforming into desired strains.

**Table 2.**
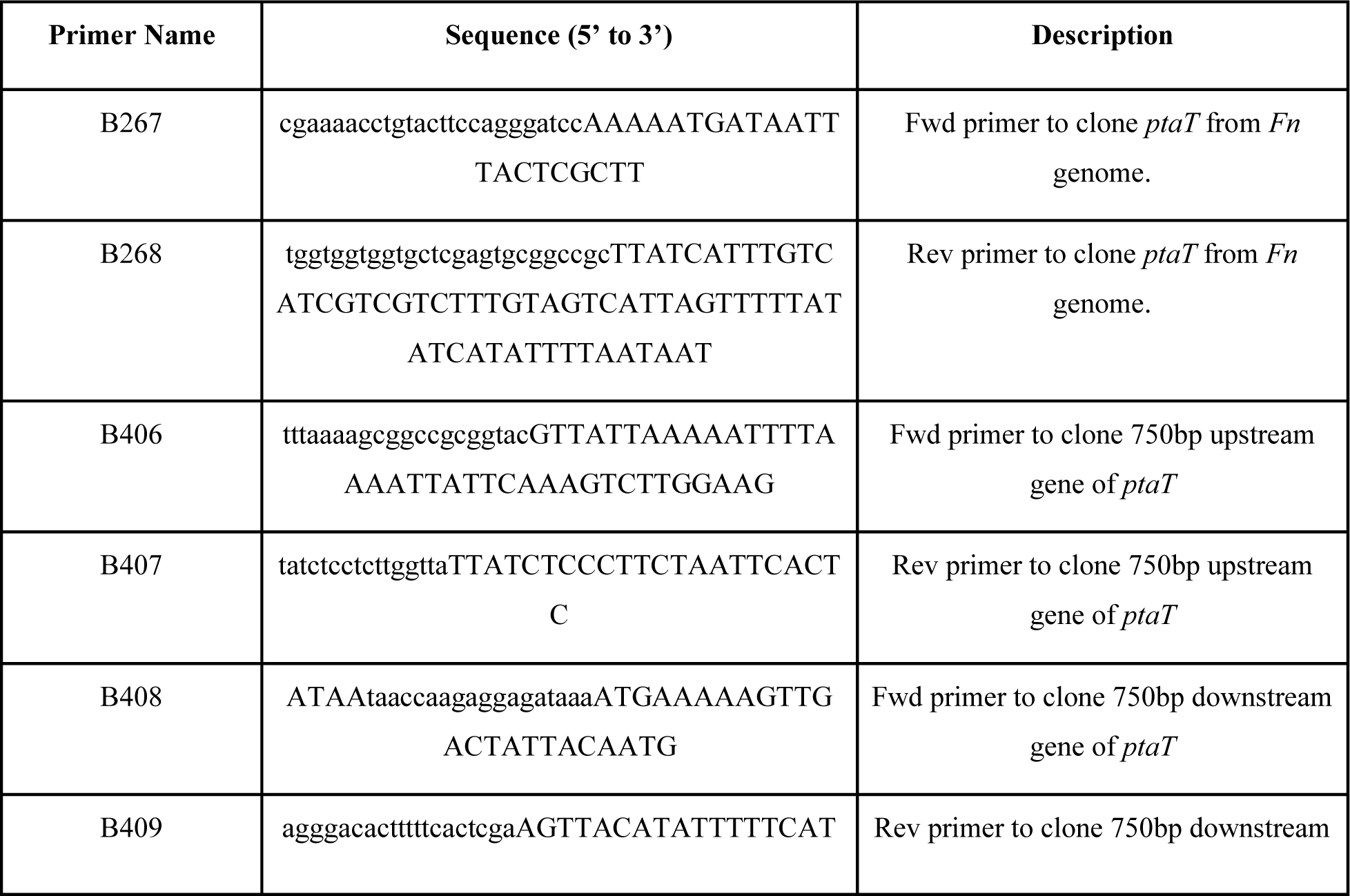
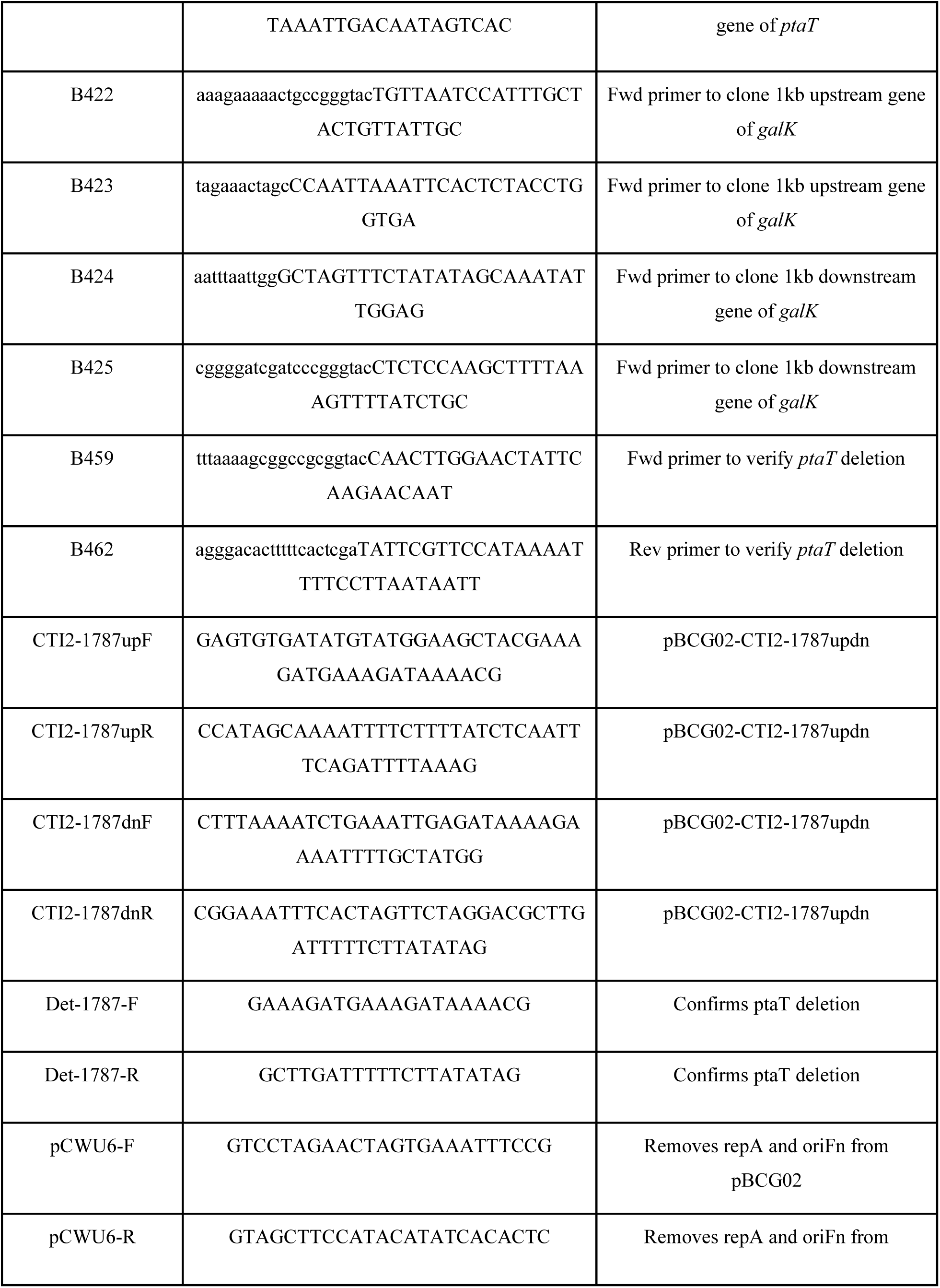
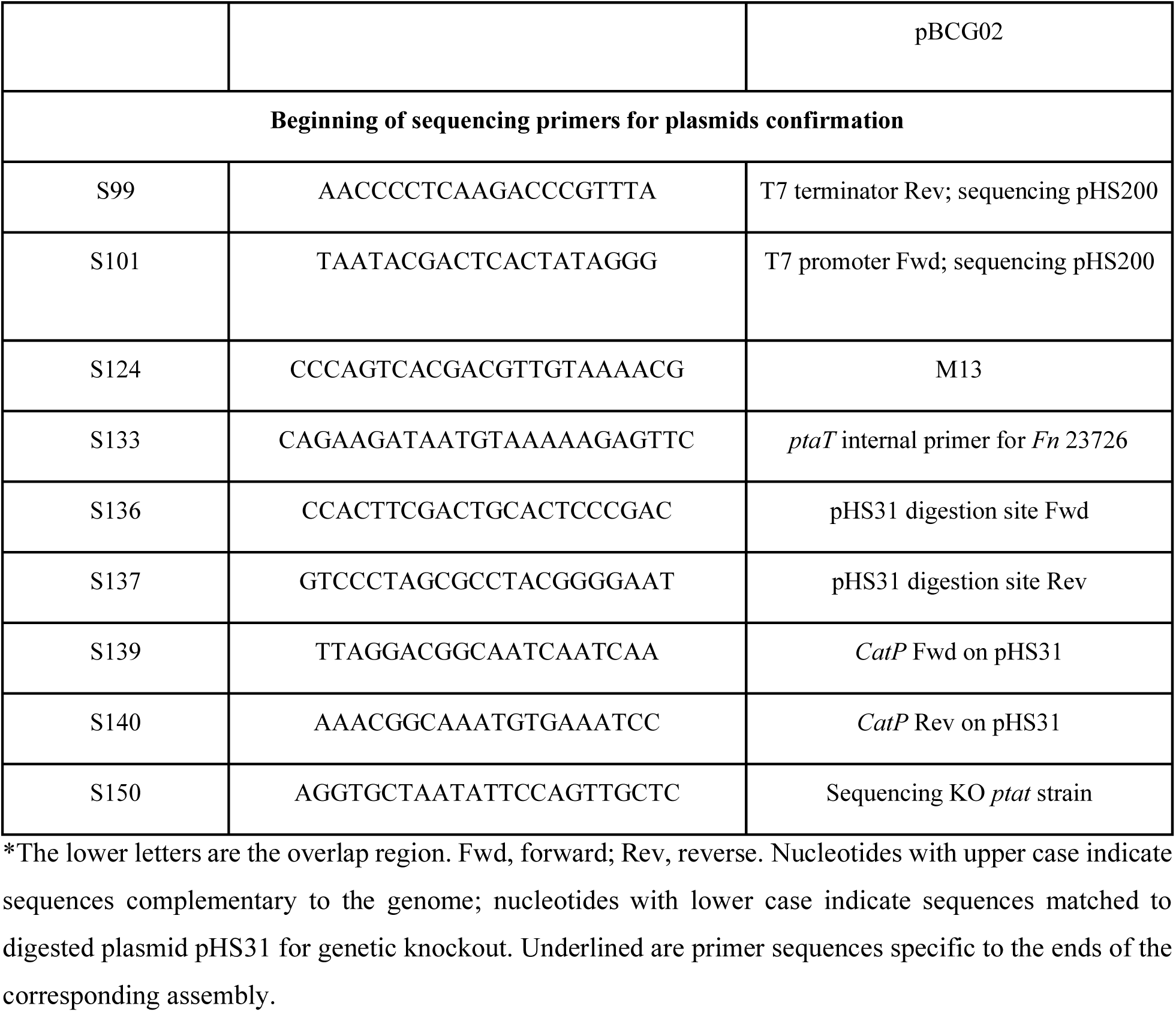
Primers used for strain construction.

### Generation of the deletion mutant in F. nucleatum ATCC 23726 via electroporation

The Δ*patT* strain was generated from the following steps. Electrocompetent *F. nucleatum* ATCC 23726 Δ*galK* was prepared by growing a 10-ml culture to log phase (OD_600_ = ∼0.8) followed by centrifugation at 12,000 × *g* for 10 min, removal of supernatant, and three successive washes with 750 μl of ice-cold electroporation buffer (10% glycerol, 1 mM MgCl_2_ in deionized H_2_O). For each electroporation, cells were resuspended in ice-cold electroporation buffer at an OD_600_ of 12. Bacteria were transferred to ice-cold 1-mm electroporation cuvettes (Fisher Scientific), and anaerobically incubated with 1 μg (concentration >500 ng/μl) of plasmid on ice for 10 min before electroporating at a setting of 2.5 kV (25 kV/cm), 25 μF and 200 Ω using a Gene Pulser Xcell Electroporation System (Bio-Rad). Immediately after electroporation, bacteria were transferred to 1 ml of anaerobically pre-reduced CB supplemented with 1mM MgCl_2_ and then incubated anaerobically at 37 °C overnight. After outgrowth, bacteria were spun down at 12,000 × *g* for 5 min, medium was removed, cells were spread on CB blood agar plates containing 5 μg/ml thiamphenicol. Plates were incubated in an anaerobic chamber at 37 °C for 3-7 days for colony growth to select for the first crossover step. Successful single crossover clones were verified by gDNA extraction, colony PCR and Sanger sequencing.

To enable the second step of crossover, sequencing verified clones were inoculated in 2 ml fresh CB broth containing 0.25% 2-deoxy-D-galactose (2-DG) and incubated under anaerobic conditions overnight at 37°C. *Fn* Δ*patT* clones were selected by spreading 100 μl of diluted culture on blood agar plate containing 0.25% 2-deoxy-D-galactose (2-DG). Plates were incubated under anaerobic conditions for 3-4 days at 37°C. 1-3 colonies were then re-streaked on the 0.25% 2-DG agar plates for another 3 days of incubation. 8-10 single colonies were randomly picked and streaked onto blood agar plates supplemented with 0.25% 2-DG. The same colonies were simultaneously streaked onto blood agar plates containing 5 μg/ml thiamphenicol to confirm thiamphenicol sensitivity as the loss of thiamphenicol resistance indicate the removal of the plasmid backbone. Subsequently, thiamphenicol-sensitive colonies were inoculated into 1% 2-DG CB broth and passaged 4 times in 1% 2-DG CB broth followed by spreading onto 0.25% 2-DG blood agar plates. 10-20 single colonies were picked, expanded for genomic DNA extraction and verified by colony PCR and Sanger sequencing to confirm the double crossover knockout for *patT*.

### Plasmid construction for genetic knockout of F. nucleatum CTI-2

PCR primers used in this study are listed in Table 2. For the deletion of CTI-2 gene *ptaT*, the plasmid pBCG02-CTI2-1787updn was constructed via Gibson assembly cloning according to the manufacturer’s instructions with NEBuilder HiFi DNA Assembly master mix. Briefly, 1.0-kb fragments of upstream and downstream regions were amplified by PCR using primer pairs CTI2-1787up-F/CTI2-1787up-R and CTI2-1787dn-F/CTI2 1787dn-R, respectively (Table 2). The overlapping PCR was used to ligate two PCR amplicons using primer pair CTI2-1787up-F/CTI2-1787dn-R. The fused segment was mixed in 10 µL Gibson assembly master mix solution with pBCG02 vector backbone generated by inverse PCR with primer pair pCWU6-F/pCWU6-R^62^. A 2 × PrimerSTAR® Max DNA Polymerase master mix from TaKaRa (catalog NO. R045A) was used for high-fidelity PCR amplification of the gene fragments and pBCG02 backbone. The resulting plasmid pBCG02-CTI2-1787updn was further confirmed by sequencing and transformed into F. nucleatum CTI-2 strain by electroporation^32^.

### Deletion of gene ptaT in F. nucleatum CTI-2 strain

Plasmid pBCG02-CTI2-1787updn intergradation into the bacterial chromosome DNA by homologous recombination was selected by anaerobic growth on TSPC agar plates with 5 µg/mL thiamphenicol at 37°C. Due to the low transformable efficiency of the CTI-2 strain, successful plasmid integration was achieved after multiple attempts. The thiamphenicol-resistant colony was next inoculated in TSPC broth without antibiotics overnight. The next day, cultures were diluted 1,000-fold, and 100-µL of aliquots were spread on TSPC agar plates supplemented with 2 mM inducer theophylline. This step was designed to select cells that had lost the plasmid via the second crossover event. Ten colonies were randomly chosen and re-streaked on TSPC plates to verify thiamphenicol sensitivity and analyzed by PCR to confirm the deletion of the target gene.

### Biotinylated tsRNA affinity pulldown from bacterial lysate

Dynabeads® M-270 Streptavidin beads (ThermoFisher, Catalog# 65305) and streptavidin agarose resin (G-Biosciences. St. Louis, MO. U.S.A) were washed according to the instructions and blocked by 1 mg/ml BSA at RT for 1h. M-270 beads and resin were washed by 3×TBS (Tris HCl 150mM, NaCl 0.45M, pH=7.4) and 1×TBS (Tris HCl 50mM, NaCl 0.15M, pH=7.4) three times respectively and resuspended in the same volume buffer as the initial volume of beads taken from vial by 3×TBS and 1×TBS. 50 ml wild type *Fn* culture was harvested by spinning at 4000 × rcf 15 min followed by 3 times 1×PBS washing steps. Cell pellet was rotated at RT for 20 min in 3 ml cell lysis buffer (50 mM Tris HCl, 0.15 M NaCl, ph7.4, 1 mM DTT, 0.5% NP-40, 50 μg/ml lysozyme and protease inhibitor cocktail EDTA free). Lysates were sonicated on ice with 5s on and 5s off for a total 5 min at 50% power (Soniprobe, Dawe Instruments, England) for 3-4 times until the lysate was transparent and cleared by centrifugation. Heparin was added to the clear extracts with 100 μg/ml working concentration followed by incubation with precoated resin at 4 °C for 30 min and quantified for protein concentration by DC Protein Assay Reagent package. Before incubating with pretreated M-270 beads, 1 mM MgCl_2_, 1 mM ATP and 40 U/ml RnaseOUT (Thermofisher) were added to the precleared cell lysates.

5’ or 3’ Biotin-tsRNA with a final concentration of 500 nM was added to 10μl of precoated M-270 beads and incubated on ice for 30 min. After 3 times of washing by 1×TBS, the precleared lysates were aliquoted to the pretreated M-270 beads equally and incubated at RT for 2h on the rotator. M-270 beads were washed for 10 min at RT with wash buffer (50mM Tris HCl, 0.15M NaCl, pH 7.4, 0.5% NP-40) twice followed by 1×TBST (25mM Tris, 0.15M NaCl, pH=7.4, 0.05% TweenTM-20) twice. 1×SDS loading buffer was added to the samples and prepared for SDS-PAGE. After SDS-PAGE, the gel was stained by FOCUS™ FASTsilver™ kit (Bioscience, 786-240). Target bands were excised and de-stained for Mass Spectrometry (Thermo Q Exactive) analysis at the Barnett Institute Proteomics Mass Spectrometry Core Facility at Northeastern University.

### Mass Spectrometry

Proteins were reduced (10 mM dithiothreitol, 56 °C for 45 min) and alkylated (50 mM iodoacetamide, room temperature in the dark for 1 h). Proteins were subsequently digested with trypsin (sequencing grade, Promega), at an enzyme/substrate ratio of 1:50, at room temperature overnight in 100 mM ammonium acetate, pH 8.9. Trypsin activity was quenched by adding formic acid to a final concentration of 5%. Peptides were desalted using C18 SpinTips (Protea) then lyophilized and stored at −80 °C. Peptides were loaded on a pre-column and separated by reverse phase HPLC (Thermo Easy nLC1000) over a 140-min gradient before nano electrospray using a QExactive Mass Spectrometer (Thermo Fisher Scientific). The Mass Spectrometer was operated in a data-dependent mode. The parameters for the full scan MS were: resolution of 70,000 across 350–2000 m/z, AGC 3e6, and maximum IT 50 ms. The full MS scan was followed by MS/MS for the top 10 precursor ions in each cycle with an NCE of 28 and dynamic exclusion of 30 s. Raw mass spectral data files (.raw) were searched using Proteome Discoverer (Thermofisher) and Mascot version 2.4.1 (Matrix Science). Mascot search parameters were: 10 ppm mass tolerance for precursor ions; 0.8 Da for fragment ion mass tolerance; 2 missed cleavages of trypsin; fixed modification was carbamidomethylation of cysteine; variable modification was methionine oxidation. Only peptides with a Mascot score greater than or equal to 25 and an isolation interference less than or equal to 30 were included in the data analysis. Potential interacting proteins are identified in the experimental sample after the removal of proteins in the control sample and common contaminating proteins.

### Recombinant protein production and purification

*E. coli* was cultured in 250 ml LB medium until OD_600_ reached ∼0.8 followed by 1mM isopropyl-β-d-thiogalactopyranoside (IPTG) induction for 18h at 20 °C. *E. coli* was harvested by centrifugation and resuspended in Binding Buffer (0.5 M NaCl, 33 mM sodium phosphate, 20 mM imidazole, pH 7.6) with 1-2 mg/ml lysozyme, 1% (v/v) Triton X-100, 1 tablet of EDTA-free Protease Inhibitor Cocktail, 1 mM PMSF, and Benzonase followed by gently rotation at RT for 20min. PMSF was re-added every 30 min until the protein was bound to Nickel resin. Lysates were sonicated on ice with 5-s on and 5-s off for a total of 5 min at 50% power (Soniprobe, Dawe Instruments, England) for 1-2 times until the lysate was not viscous and cleared by centrifugation for 1h at 12000×rcf, 4°C. 1ml Nickel resin was washed by deionized H_2_O two times followed by the binding buffer washing. After the sonicated media was fully centrifuged, the supernatant was poured into 15-ml conical tube with Nickel resin and 1% Triton-114 was added to the supernatant. After 1h incubation at 4°C, the resin was washed via the binding buffer containing 1% Triton-114 three times at 4°C, 30 min per wash. Resin was transferred to the polypropylene column (Bio-Rad) and eluted with Elution Buffer (0.5 M NaCl, 33 mM sodium phosphate, 250 mM imidazole, pH 7.6). The protein elute was concentrated to ∼ 0.5 ml by 10kDa MWCO Spin Column, and then loaded onto a size exclusion column, ENrich™ SEC 650 10 x 300 (Bio-Rad), which was connected to the NGC Medium-Pressure Liquid Chromatography System (Bio-Rad). The protein concentration was monitored with OD_280_, and a fraction collector (Bio-Rad) was used to collect samples at 0.5 ml per fraction for SDS-PAGE analyses. Only fractions containing the desired protein without any impurity were pooled for downstream binding assays. After FPLC, the protein samples were added with 1mM DTT, and stored at -80°C before use.

### Direct binding assay for tsRNA and purified proteins

Dynabeads® M-270 Streptavidin beads were prepared according to the manual. The concentration of purified proteins was measured by Pierce™ Rapid Gold BCA Protein Assay Kit. Purified PtaT was adjusted to 1μg per reaction by adjusting buffer (50mM Tris HCl, 0.15 M NaCl, pH 7.4, 0.1% NP-40, 1 mM ATP, 1 mM MgCI_2_, 100 ug/ml heparin,1mg/ml BSA) and incubated with biotinylated tsRNA pretreated beads at RT for 2h followed by washing steps as above described. 2 x SDS loading buffer was added to washed beads for Western Blot analysis.

### Fluorescence microscopy of Cy3 tsRNA labeled Fn strains

Cy3 labeled tsRNA was reconstituted in 1×TBS containing 0.1 mM EDTA for imaging. Overnight-grown *Fn* strains was diluted to OD_600_=0.1 and treated with 500 nM of Cy3 labeled tsRNA-000794, and scrambled RNA for 5 hours at an anaerobic chamber. Labeled bacteria were then washed with 1×PBS for three times under a centrifuge speed of 17,000×g for 10 min. Washed samples were then sandwiched between a cover glass and poly-L-lysine coated cover slide. Samples were then immediately imaged by a ZEISS LSM 800 confocal microscope with a fast Airyscan detector (with 120-nm lateral resolution and 350-nm axial resolution). To ensure the image quality, we utilized a 63× Plan-Apochromat NA=1.4 oil immersion objective. Samples were excited at a wavelength of 514 nm with a 10% power and detected in the range of 550-600 nm. To quantify the fluorescence intensity from the same sample patch, dynamic range was adjusted to be the same under a channel-mode confocal modality. To have a clear visualization of Cy3-tsRNA incorporation at subcellular level, super-resolution by Airyscan was achieved at a gain of 800 V. Images were visualized and analyzed by FiJi (NIH) and quantitative analysis was conducted by R.

### Western Blotting

The samples were first separated by 10% SDS-PAGE gel and then transferred to a nitrocellulose membrane (Fisher Scientific). The membranes were incubated with the Anti-FLAG epitope (DYKDDDDK, Biolegend, San Diego, CA, catalog# 637301) with 1:2000 dilution overnight in the cold room, and the secondary antibody anti-rat IgG HRP (Cell Signaling Technology, Danvers, MA, catalog# 7077) with the same dilution at room temperature for 1 hr. Premixed Pierce^TM^ 3,3 diaminobenzidine (DAB) substrate (Fisher Scientific) was used to detect the target proteins.

### Statistical analysis

All statistical analyses and quantitative graphs were performed using R. Statistical significance was achieved through Student’s unpaired *t*-test: ***, *p* < 0.001; **, *p* < 0.01; *, *p* < 0.05.

## Acknowledgments

This work was supported by NSF 2333230 (JL), NIH National Institute of Dental and Craniofacial Research (NIDCR) awards, DE030943 (XH), DE023810 (XH), DE029479 (XH) and DE031329 (JL), DE030895 (CW), T90 DE026110 and K99 DE033794 (to P.-T.D.). We are grateful to Dr. Susan E. Abbatiello, the director of the Barnett Institute of Chemical and Biological Analysis, Department of Chemistry and Chemical Biology at Northeastern University for scientific advice.

## REFERENCES

1 Chaplin, D. D. Overview of the immune response. Journal of Allergy and Clinical Immunology 125, S3–S23 (2010).

2 Zheng, D., Liwinski, T. & Elinav, E. Interaction between microbiota and immunity in health and disease. Cell Research 30, 492–506 (2020).

3 Belkaid, Y. & Hand, Timothy W. Role of the microbiota in immunity and inflammation. Cell 157, 121–141 (2014).

4 Malard, F., Dore, J., Gaugler, B. & Mohty, M. Introduction to host microbiome symbiosis in health and disease. Mucosal Immunology 14, 547–554 (2021).

5 Li, M. et al. Symbiotic gut microbes modulate human metabolic phenotypes. Proceedings of the National Academy of Sciences 105, 2117–2122 (2008).

6 Katiyar-Agarwal, S. & Jin, H. Role of small RNAs in host-microbe interactions. Annual Review of Phytopathology 48, 225–246 (2010).

7 Chandan, K., Gupta, M. & Sarwat, M. Role of host and pathogen-derived microRNAs in immune regulation during infectious and inflammatory diseases. Frontiers in Immunology 10 (2020).

8 Huang, C.-Y., Wang, H., Hu, P., Hamby, R. & Jin, H. Small RNAs – Big players in plant-microbe interactions. Cell Host & Microbe 26, 173–182 (2019).

9 Ahmadi Badi, S., et al. Small RNAs in outer membrane vesicles and their function in host-microbe interactions. Frontiers in Microbiology 11 (2020).

10 Koeppen, K. et al. A novel mechanism of host-pathogen interaction through sRNA in bacterial outer membrane vesicles. PLOS Pathogens 12, e1005672 (2016).

11 Choi, J.-W., Kim, S.-C., Hong, S.-H. & Lee, H.-J. Secretable small RNAs via outer membrane vesicles in periodontal pathogens. Journal of Dental Research 96, 458–466 (2017).

12 Zhang, L., Liu, J. & Hou, Y. Classification, function, and advances in tsRNA in non-neoplastic diseases. Cell Death & Disease 14, 748 (2023).

13 Kim, H. K., Yeom, J.-H. & Kay, M. A. Transfer RNA-derived small RNAs: Another layer of gene regulation and novel targets for disease therapeutics. Molecular Therapy 28, 2340–2357 (2020).

14 Liu, B. et al. Deciphering the tRNA-derived small RNAs: origin, development, and future. Cell Death & Disease 13, 24 (2021).

15 Krishna, S. et al. Dynamic expression of tRNA-derived small RNAs define cellular states. EMBO reports 20, e47789 (2019).

16 He, X. et al. Human tRNA-derived small RNAs modulate host–oral microbial interactions. Journal of Dental Research 97, 1236–1243 (2018).

17 Pandey, K. K. et al. Regulatory roles of tRNA-derived RNA fragments in human pathophysiology. Molecular Therapy - Nucleic Acids 26, 161–173 (2021).

18 Han, Y. W. *Fusobacterium nucleatum*: a commensal-turned pathogen. Current Opinion in Microbiology 23, 141–147 (2015).

19 Brennan, C. A. & Garrett, W. S. *Fusobacterium nucleatum* — symbiont, opportunist and oncobacterium. Nature Reviews Microbiology 17, 156–166 (2019).

20 Wang, N. & Fang, J.-Y. *Fusobacterium nucleatum*, a key pathogenic factor and microbial biomarker for colorectal cancer. Trends in Microbiology 31, 159–172 (2023).

21 Hong, M., et al. *Fusobacterium nucleatum* aggravates rheumatoid arthritis through FadA-containing outer membrane vesicles. Cell Host & Microbe 31, 798–810.e797 (2023).

22 Signat, B., Roques, C., Poulet, P. & Duffaut, D. Role of *Fusobacterium nucleatum* in periodontal health and disease. Current Issues in Molecular Biology 13, 25–36 (2011).

23 Liu, P. et al. Detection of Fusobacterium Nucleatum and fadA Adhesin Gene in Patients with Orthodontic Gingivitis and Non-Orthodontic Periodontal Inflammation. PLOS ONE 9, e85280 (2014).

24 Bullman, S. et al. Analysis of *Fusobacterium* persistence and antibiotic response in colorectal cancer. Science 358, 1443–1448 (2017).

25 Kostic, Aleksandar D., et al. *Fusobacterium nucleatum* potentiates intestinal tumorigenesis and modulates the tumor-immune microenvironment. Cell Host & Microbe 14, 207–215 (2013).

26 Yang, M. et al. Targeting *Fusobacterium nucleatum* through chemical modifications of host-derived transfer RNA fragments. The ISME Journal 17, 880–890 (2023).

27 Jazurek, M., Ciesiolka, A., Starega-Roslan, J., Bilinska, K. & Krzyzosiak, W. J. Identifying proteins that bind to specific RNAs -focus on simple repeat expansion diseases. Nucleic Acids Research 44, 9050–9070 (2016).

28 Abramson, J. et al. Accurate structure prediction of biomolecular interactions with AlphaFold 3. Nature (2024).

29 Kühlbrandt, W. Biology, structure and mechanism of P-type ATPases. Nature Reviews Molecular Cell Biology 5, 282–295 (2004).

30 Salustros, N. et al. Structural basis of ion uptake in copper-transporting P_1B_-type ATPases. Nature Communications 13, 5121791 (2022).

31 Tsuda, T. & Toyoshima, C. Nucleotide recognition by CopA, a Cu^+^-transporting P-type ATPase. The EMBO Journal 28, 1782–1791 (2009).

32 Peluso, E. A., Scheible, M., Ton-That, H. & Wu, C. Genetic manipulation and virulence assessment of *Fusobacterium nucleatum*. Current Protocols in Microbiology 57, e104 (2020).

33 Casasanta, M. A., et al. *Fusobacterium nucleatum* host-cell binding and invasion induces IL-8 and CXCL1 secretion that drives colorectal cancer cell migration. Science Signaling 13, eaba9157 (2020).

34 Abranches, J., Chen, Y.-Y. M. & Burne, R. A. Galactose metabolism by *Streptococcus mutans*. Applied and Environmental Microbiology 70, 6047–6052 (2004).

35 Tian, J. et al. Acquisition of the arginine deiminase system benefits epiparasitic Saccharibacteria and their host bacteria in a mammalian niche environment. Proceedings of the National Academy of Sciences 119, e2114909119 (2022).

36 Jaishankar, J. & Srivastava, P. Molecular basis of stationary phase survival and applications. Frontiers in Microbiology 8 (2017).

37 Veselovsky, V. A. et al. The gene expression profile differs in growth phases of the *Bifidobacterium* longum culture. Microorganisms 10, 1683791 (2022).

38 Bathke, J., Konzer, A., Remes, B., McIntosh, M. & Klug, G. Comparative analyses of the variation of the transcriptome and proteome of Rhodobacter sphaeroides throughout growth. BMC Genomics 20, 358 (2019).

39 Oba, Y. et al. Identifying the wide diversity of extraterrestrial purine and pyrimidine nucleobases in carbonaceous meteorites. Nature Communications 13, 2008791 (2022).

40 Egli, M. & Manoharan, M. Chemistry, structure and function of approved oligonucleotide therapeutics. Nucleic Acids Research 51, 2529–2573 (2023).

41 Cheng, J.-X. & Xie, X. S. Vibrational spectroscopic imaging of living systems: An emerging platform for biology and medicine. Science 350, aaa8870 (2015).

42 Butler, H. J. et al. Using Raman spectroscopy to characterize biological materials. Nature Protocols 11, 664–687 (2016).

43 Dong, P.-T. et al. Polarization-sensitive stimulated Raman scattering imaging resolves amphotericin B orientation in *Candida* membrane. Science Advances 7, eabd5230 (2021).

44 Gardner-Lubbe, S. Linear discriminant analysis for multiple functional data analysis. Journal of Applied Statistics 48, 1917–1933 (2021).

45 Xanthopoulos, P., Pardalos, P. M. & Trafalis, T. B. in Robust Data Mining 27–33 (Springer New York, 2013).

46 Xu, J. et al. Artificial intelligence-aided rapid and accurate identification of clinical fungal infections by single-cell Raman spectroscopy. Frontiers in Microbiology 14 (2023).

47 Ember, K. J. I. et al. Raman spectroscopy and regenerative medicine: a review. npj Regenerative Medicine 2, 12 (2017).

48 Rubinstein, M. R., et al. *Fusobacterium nucleatum* promotes colorectal cancer by inducing Wnt/catenin modulator Annexin A1. EMBO reports 20, e47638 (2019).

49 Abed, J. et al. Fap2 mediates *Fusobacterium nucleatum* colorectal adenocarcinoma enrichment by binding to tumor-expressed Gal-GalNAc. Cell Host & Microbe 20, 215–225 (2016).

50 Dar, D. & Sorek, R. Bacterial noncoding RNAs excised from within protein-coding transcripts. mBio 9, 10.1128/mbio.01730-01718 (2018).

51 Lambert, M., Benmoussa, A. & Provost, P. Small Non-Coding RNAs Derived from Eukaryotic Ribosomal RNA. Non-Coding RNA 5, 16 (2019).

52 Yamamura, S., Imai-Sumida, M., Tanaka, Y. & Dahiya, R. Interaction and cross-talk between non-coding RNAs. Cellular and Molecular Life Sciences 75, 467–484 (2018).

53 Liu, S. et al. The host shapes the gut microbiota via fecal microRNA. Cell Host & Microbe 19, 32–43 (2016).

54 Li, M., Chen, W.-D. & Wang, Y.-D. The roles of the gut microbiota–miRNA interaction in the host pathophysiology. Molecular Medicine 26, 101 (2020).

55 Feinberg, E. H. & Hunter, C. P. Transport of dsRNA into cells by the transmembrane protein SID-1. Science 301, 1545–1547 (2003).

56 Chen, Q. et al. SIDT1-dependent absorption in the stomach mediates host uptake of dietary and orally administered microRNAs. Cell Research 31, 247–258 (2021).

57 Nguyen, T. A. et al. SIDT2 transports extracellular dsRNA into the cytoplasm for innate immune recognition. Immunity 47, 498–509. e496 (2017).

58 Chan, H. et al. The P-Type ATPase superfamily. Journal of Molecular Microbiology and Biotechnology 19, 5–104 (2010).

59 Palmgren, M. P-type ATPases: Many more enigmas left to solve. Journal of Biological Chemistry 299 (2023).

60 Fujii, T. et al. Parkinson’s disease-associated ATP13A2/PARK9 functions as a lysosomal H^+^,K^+^-ATPase. Nature Communications 14, 2174791 (2023).

61 Mu, J. et al. Conformational cycle of human polyamine transporter ATP13A2. Nature Communications 14, 1978791 (2023).

62 Gc, B., Zhou, P. & Wu, C. HicA toxin-based counterselection marker for allelic exchange mutations in *Fusobacterium nucleatum*. Applied and Environmental Microbiology 89, e00091–00023 (2023).

