## Supplementary information for "Identification of P-type ATPase as a bacterial transporter for host-derived small RNA"

This document includes:

Figure S1-S9

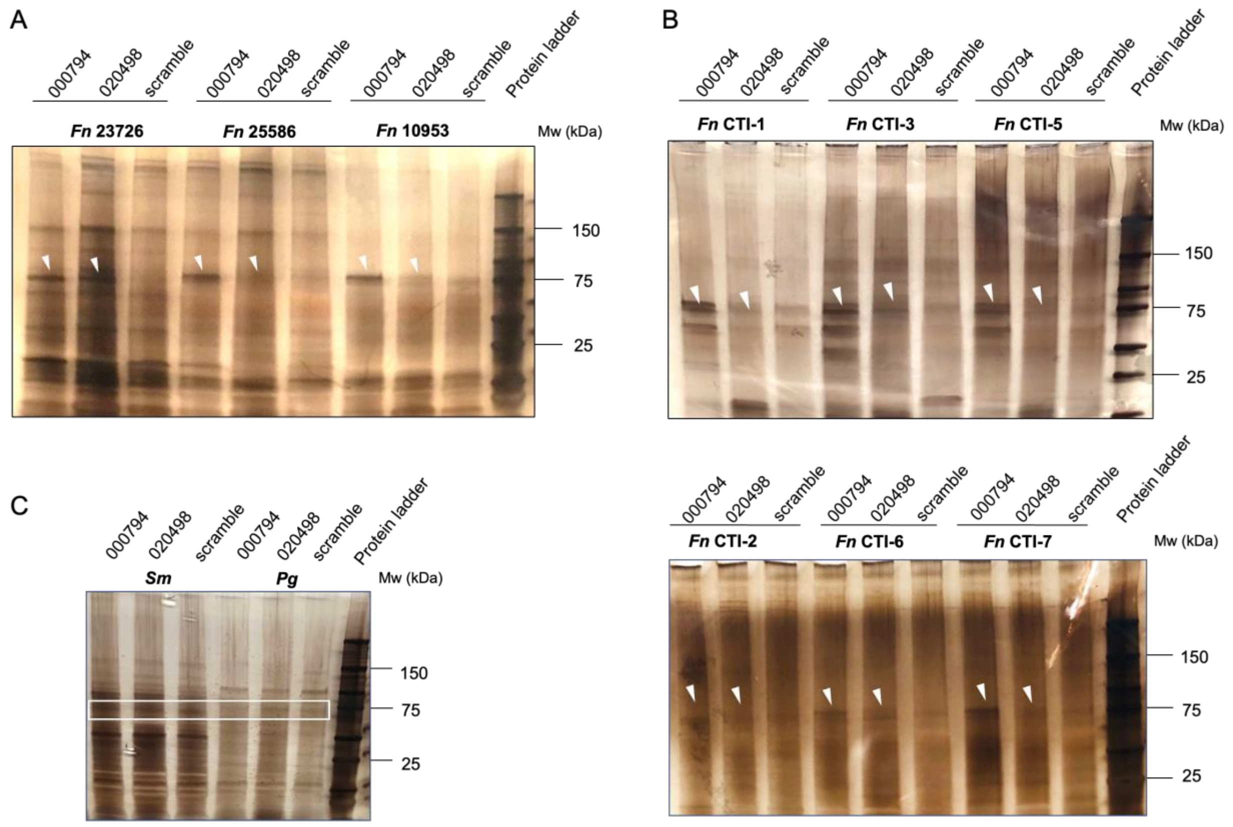


**Supplementary Figure 1. Identification of PtaT by tsRNA-mediated affinity pulldown. A.** Silver staining of denaturing SDS-PAGE gel for biotinylated tsRNA pulldown samples in three different ATCC *Fn* strains. Arrowheads indicate the gel bands, which were excised for protein identification by Mass Spectrometry analysis. **B.** Silver staining of denaturing SDS-PAGE gel for biotinylated tsRNA pulldown samples in six *Fn* clinical tumor isolates (CTIs). Arrowheads indicate the gel bands, which were excised for protein identification by Mass Spectrometry analysis. CTI-1: *F. nucleatum. ssp. animalis*; CTI-3: *F. nucleatum. ssp. animalis*; CTI-5: *F. nucleatum. ssp. animalis*; CTI-2: *F. nucleatum, ssp. nucleatum*; CTI-6: *F. nucleatum. ssp. polymorphum*; CTI-7: *F. nucleatum. ssp. vincentii.* **C.** Silver staining of denaturing SDS-PAGE gel for biotinylated tsRNA pulldown samples in *Streptococcus mitis ATCC 6249* (*Sm*) and *Porphyromonas gingivalis ATCC 33277* (*Pg*).

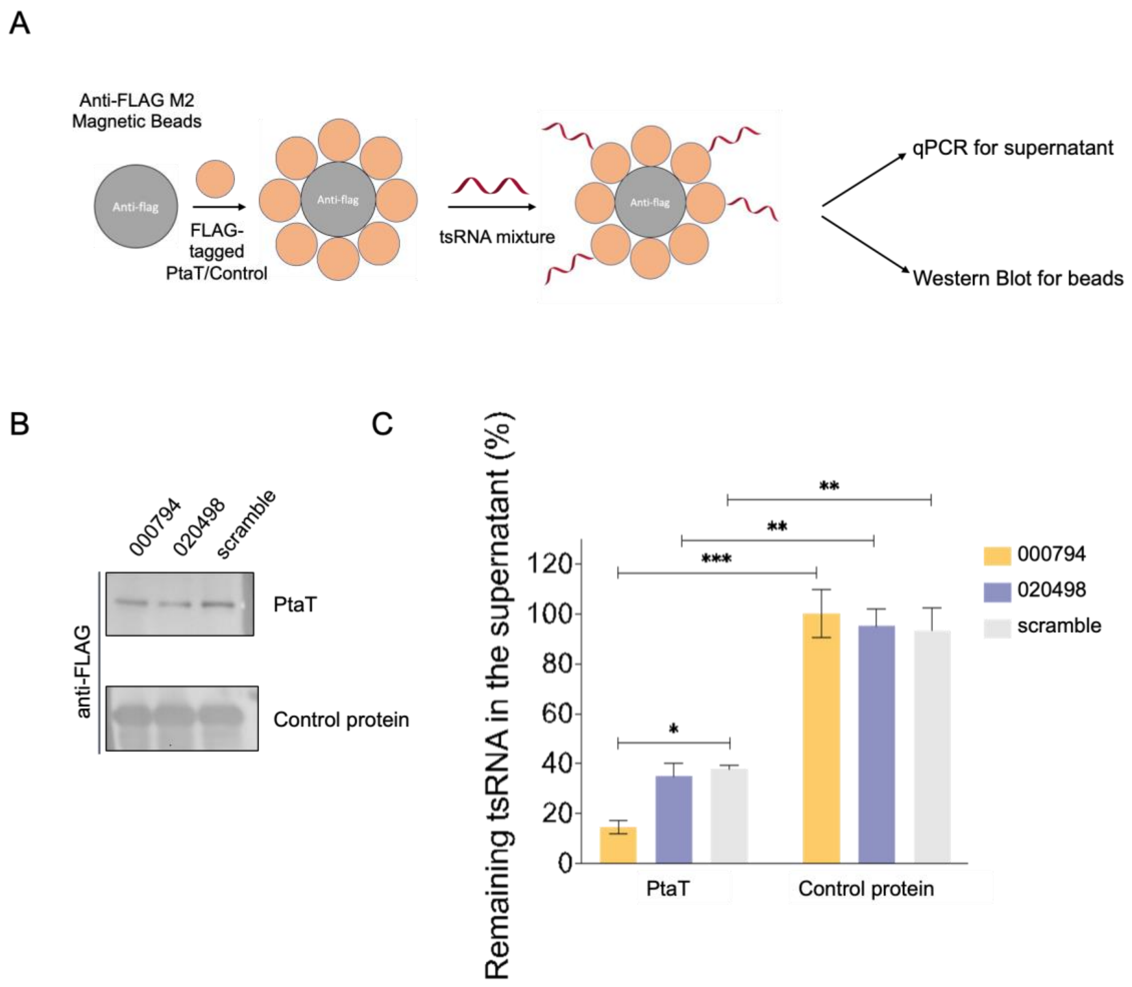


**Supplementary Figure 2. Validation of the binding interaction between PtaT and tsRNA. A.** Schematic of using anti-FLAG M2 magnetic beads and FLAG-tagged recombinant proteins to validate tsRNA binding *in vitro*. **B.** Western blotting of FLAG-tagged PtaT and STING (a negative control known to bind cyclic dinucleotides but not RNA oligos). **C.** Pull-down of naturally occurring tsRNA-000794, tsRNA-020498 or scramble control by purified FLAG-tagged PtaT and anti-FLAG antibody-conjugated magnetic beads. A FLAG-tagged irrelevant protein (STING) was used as a negative control for nonspecific binding. Unbound RNAs were quantified by stem-loop qPCR and normalized to the initial concentration. Results = Mean ± SEM (N=4) and are representative of two biological replicates. Statistical analyses were performed by the two-way ANOVA followed by Dunnett's Bonferroni multiple comparison tests. *$p < 0.05$, ** $p < 0.01$, *** $p < 0.001$.

| Classification | Protein | Localization | Uniport | Groups |
| --- | --- | --- | --- | --- |
| RNA metabolism | Polyribonucleotide nucleotidyltransferase | Cytoplasmic | D5RFI6 | |
| Metal ion transport | Heavy metal translocating P-type ATPase | Membrane | D5RD38 | **tsRNA000794** |
| | Uncharacterized | | D5RA61 | |
| Metabolic process | S-methyl-5-thioribose-1-phosphate isomerase | | D5RDB1 | |
| Metabolic process | Signal peptide peptidase SppA | Membrane | D5REW1 | |
| Transport | ABC transporter, substrate-binding protein | | D5RBV3 | **tsDNA000794** |
| | Uncharacterized | | D5RA61 | |
| RNA metabolism | Ribonuclease J | Cytoplasmic | D5RD58 | |
| RNA metabolism | Polyribonucleotide nucleotidyltransferase | Cytoplasmic | D5RFI6 | **piRNA_016792** |
| | Uncharacterized | | D5RA61 | |
| RNA metabolism | Polyribonucleotide nucleotidyltransferase | Cytoplasmic | D5RFI6 | |
| RNA metabolism | Ribonuclease R | Cytoplasmic | D5RAL6 | **piRNA_006465** |
| | Uncharacterized | | D5RA61 | |
| Metabolic process | S-methyl-5-thioribose-1-phosphate isomerase | | D5RDB1 | **Beads only** |

**Supplementary Figure 3. Affinity pulldown from *Fn* ATCC 23726 total lysate examined by Mass Spectrometry.**

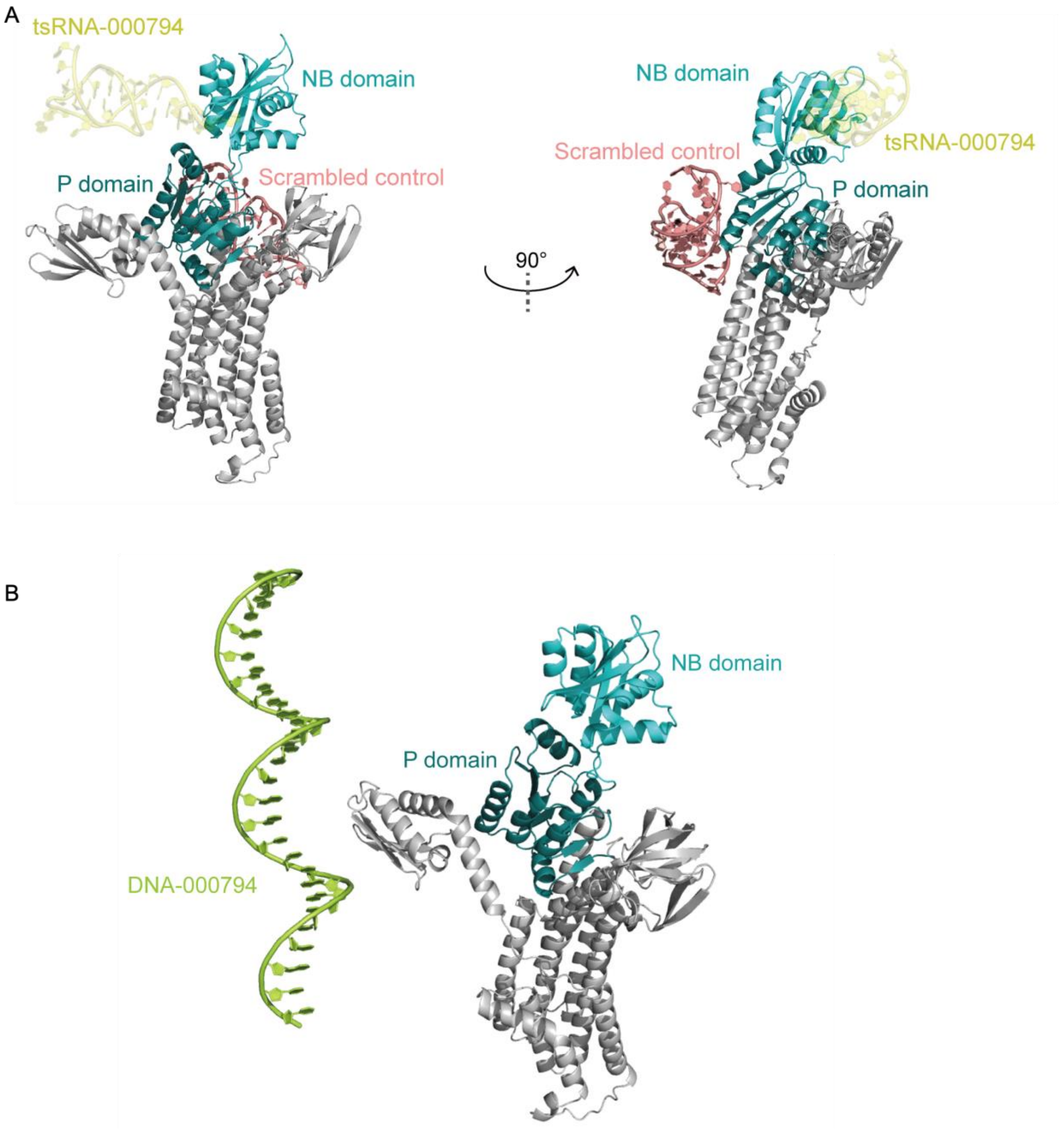


**Supplementary Figure 4. Predication of the interaction between scrambled control (A), DNA-000794 (B) and PtaT by AlphaFold 3.**

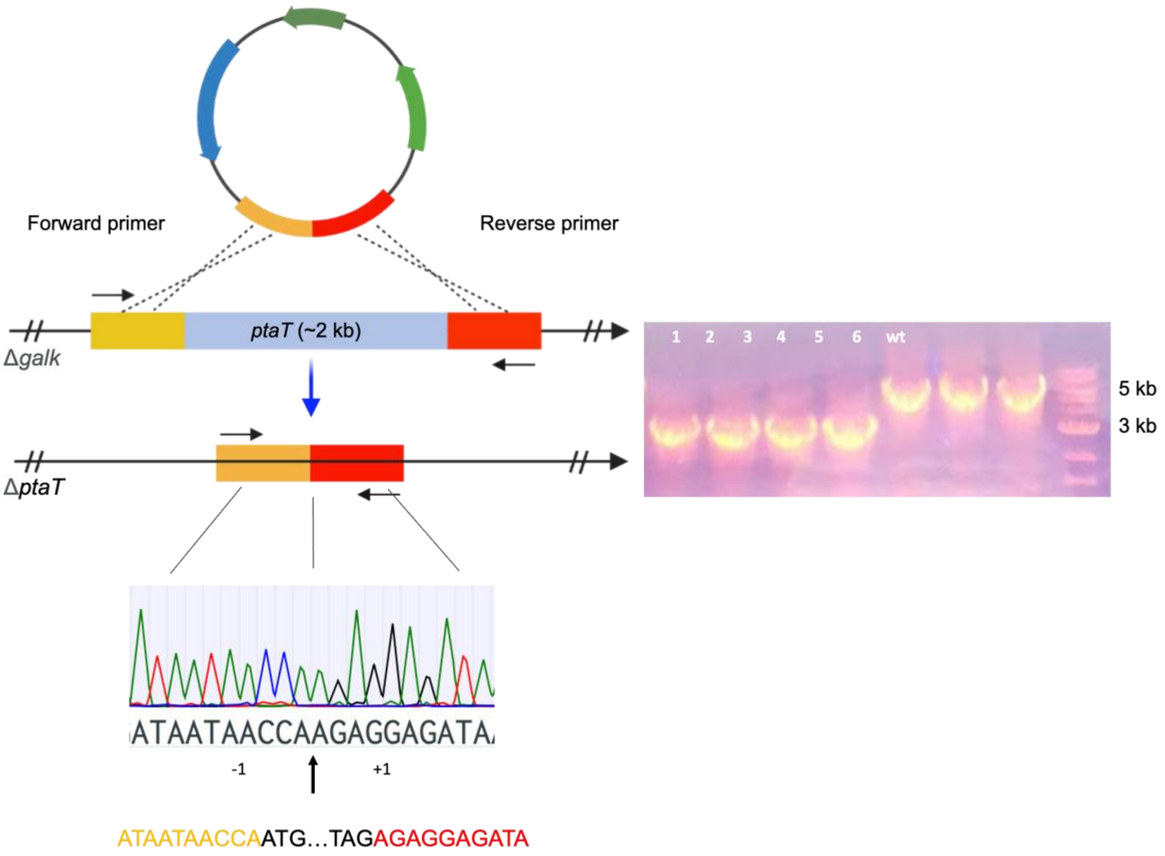


**Supplementary Figure 5. Generation of insertional mutagenesis via a double crossover-mediated complete knockout of the full-length *ptaT* in *Fn* Δ*galK*.**

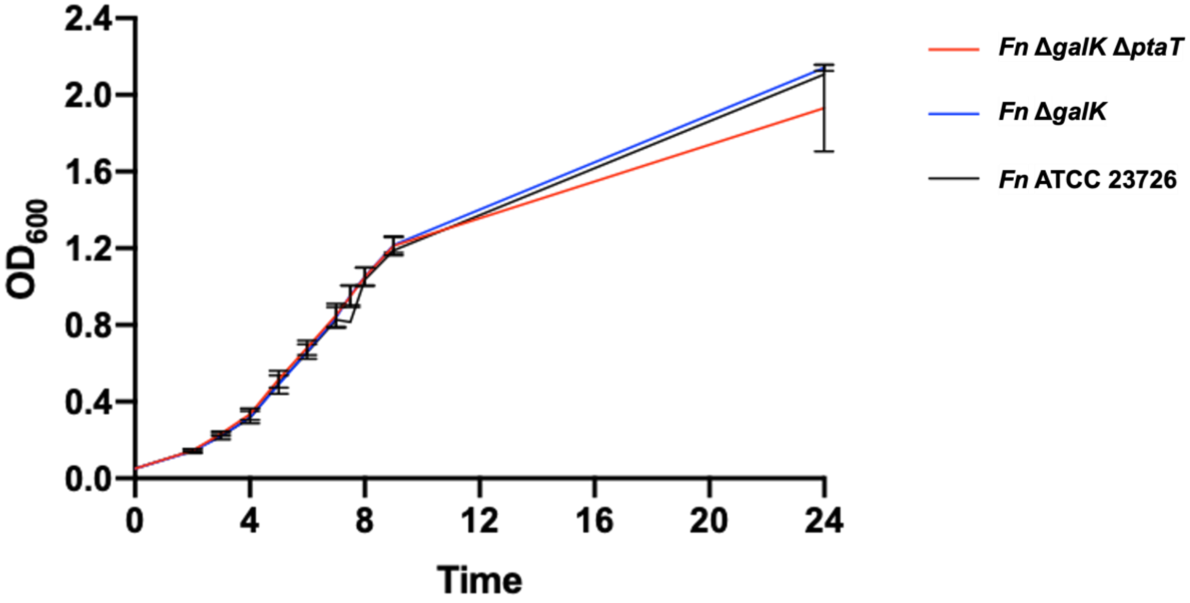


**Supplementary Figure 6**. **The time-course growth kinetic monitored by optical density at 600 nm ($OD_{600}$) from *Fn* ATCC 23726, *Fn* Δ*galK* and *Fn* Δ*galK* Δ*ptaT*.**

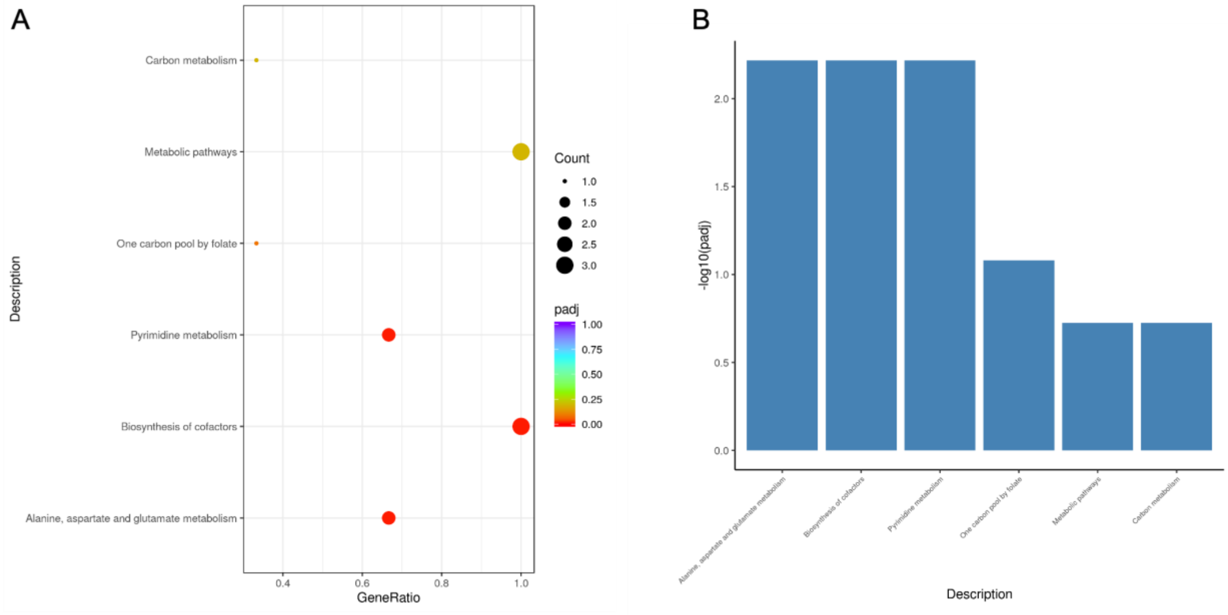


**Supplementary Figure 7. Clusters of orthologous groups (COG, A) and quantification of differentially expressed genes (B) from stationary-phase *Fn* Δ*galK* Δ*ptaT* relative to *Fn* Δ*galK*.**

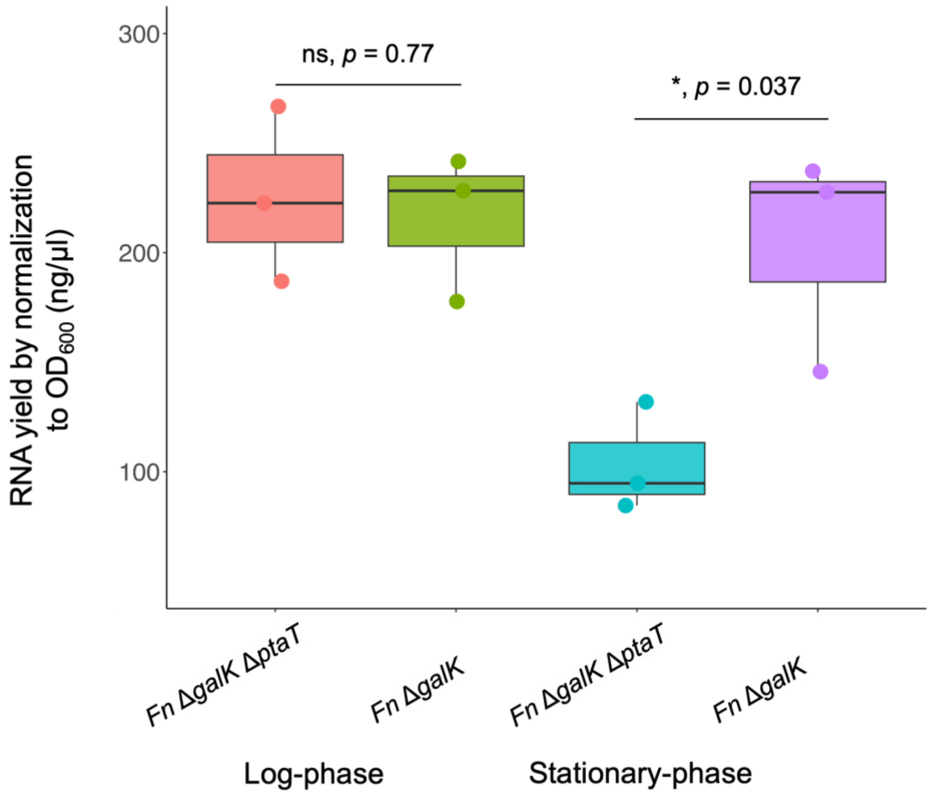


**Supplementary Figure 8. The yield of total extracted RNA of both *Fn ΔgalK* and *Fn ΔgalK ΔptaT* from three biological replicates.**

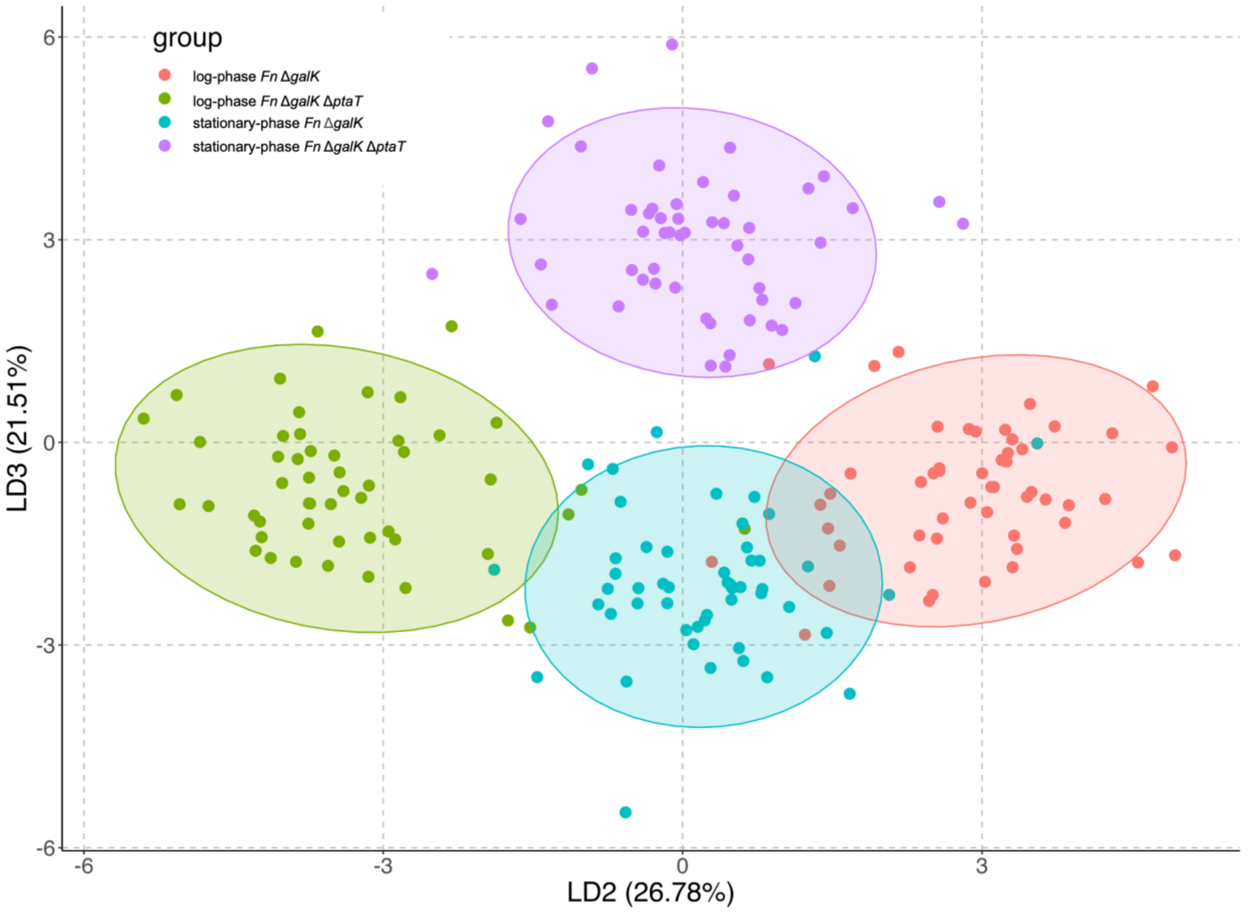


**Supplementary Figure 9. LDA analysis (LD3 versus LD2) of 200 Raman spectra from log-phase and stationary-phase *Fn ΔgalK* and *Fn ΔgalK ΔptaT*.**
